# Postmitotic Regulation of Neuronal Subpopulations in the Cerebellum

**DOI:** 10.64898/2026.08.09.743777

**Authors:** Pamela Valnegri, Shin-ichiro Fujita, Tomoko Yamada, Yue Yang

## Abstract

Hundreds of neuronal cell types and subtypes have been identified through transcriptome profiling of the mammalian brain and are thought to arise from lineage-restricted neural progenitors during early development. However, how neurons might further diversify their transcriptional identities during postmitotic development and through adulthood remains poorly understood. Here, we combine genetic and biochemical approaches to uncover granule neuron subpopulations in the anterior cerebellum, which we term C1 and C2. We find that a large proportion of granule neurons predominantly express the C2 gene program during early postmitotic differentiation, but that C2 genes are downregulated in a subset of neurons during late postnatal development, leading to comparable proportions of C1 and C2 neurons in adulthood. Using an *in vivo* genetic mini-screen, we identify calcium signaling pathways, together with the transcription factor ETV1, that establish the transcriptional program of C2 neurons. Finally, C1 and C2 granule neurons are differentially engaged during behavior and this reflects their roles in cerebellar-dependent associative learning. Together, these findings reveal postmitotic mechanisms that continue to shape neuronal identity in the brain.

## Introduction

The brain comprises diverse neuronal cell types that assemble into specialized neural circuits to support sensory, motor, and cognitive functions. During early brain development, neural precursors divide to expand the brain, giving rise to lineage-restricted mitotic progenitors and distinct postmitotic cell types. A key theme that has emerged is that the timing and location of neural precursor divisions influence the cell types produced^1–5^. In addition, advances in single cell transcriptome profiling and lineage tracing techniques have revealed that, following neuronal specification, immature postmitotic neurons may further differentiate into distinct subtypes^6–9^. However, whether mature neurons, after integrating into neural circuits, continue to diversify their transcriptional identities into adulthood remains unclear.

Cerebellar granule neurons are the most abundant neurons in the mammalian brain and provide an ideal system to study the molecular mechanisms that control neuronal development and plasticity^10, 11^. Transcriptome profiling has identified gene expression patterns that define granule neuron subtypes in anterior, posterior, and flocculonodular lobes of the cerebellum^12^, potentially reflecting their anatomical connections and roles in behavior^13–15^. However, it remains unknown whether granule neurons within individual cerebellar regions can be transcriptionally distinguished, and if so, what mechanisms regulate this process.

Neurons respond to cues from their local cellular niche as well as from long-range synaptic inputs. These cues activate transmembrane receptors and intracellular signaling cascades that may engage transcription factors in the nucleus to initiate gene expression programs. For example, the activation of NMDA receptors facilitates calcium influx, which leads to the phosphorylation of transcription factors and induces activity-dependent genes associated with neural circuit plasticity and memory^16^. Although neurons express numerous receptors responsive to neurotransmitters, neuromodulators, and growth factors, we still lack resolution into the specific roles and interactions of these pathways in regulating gene transcription and neuronal identity, particularly in the brain *in vivo*.

In this study, we characterized the molecular identities of granule neurons in the anterior cerebellum, where we recently identified neural circuits important for sensorimotor learning. We found that the anterior cerebellum contained transcriptionally distinguishable granule neuron subpopulations, termed C1 and C2, which are defined by hundreds of genes and are partially separated within the internal granule layer (IGL) of the cerebellar cortex. During early postmitotic development through synaptogenesis, the C2 gene expression program was robustly upregulated, which led to a predominance of granule neurons that preferentially expressed the C2 program. Later, from the juvenile to the young adult stage, the C2 program was downregulated in a subset of granule neurons, increasing the proportion of granule neurons preferentially expressing the C1 program. Using an *in vivo* genetic mini-screen combined with RNA-seq and ChIP-seq analyses, we identified the transcription factor ETV1 and transmembrane regulators of calcium signaling as key drivers of C2 gene expression and regulatory enhancer activity. In behaving mice, C2 granule neurons exhibited robust induction of activity-dependent transcriptional responses during high locomotor activity, but showed weaker induction than C1 granule neurons during low locomotor activity. Consistent with this activity pattern, C2 granule neurons induced cerebellar-dependent associative learning selectively during locomotion but not while mice were at rest. Together, our study reveals transcriptionally distinguishable subpopulations of postmitotic granule neurons, mechanisms that contribute to their identity, and their functions in sensorimotor learning.

## Results

### Identification of granule neuron subpopulations in the anterior cerebellum

We previously identified roles for the anterior dorsal cerebellar vermis (ADCV), the dorsal part of lobule IV/V, in associative sensorimotor learning in mice^10^. We first characterized the gene expression profiles of granule neurons in this brain region by examining published <u>s</u>ingle <u>n</u>ucleus RNA-seq (snRNA-seq) data from cerebellar lobule IV/V of adult mice (Fig. 1a)^12^ and performed unsupervised clustering to partition granule neurons into two major subpopulations (Fig. 1b; Methods). These subpopulations were defined by the differential expression of hundreds of genes, including *Syt1* and *Cadm1* enriched in cluster 1 (C1) and *Syt2* and *Cntnap4* enriched in cluster 2 (C2; Fig. 1b; Supplementary Table 1). We also observed granule neurons with intermediate C1 and C2 expression profiles (Fig. 1c; Extended Data Fig. 1a, b), consistent with findings of continuous transcriptional variation among neuronal subtypes in other brain regions^17–19^.

**Fig. 1.**
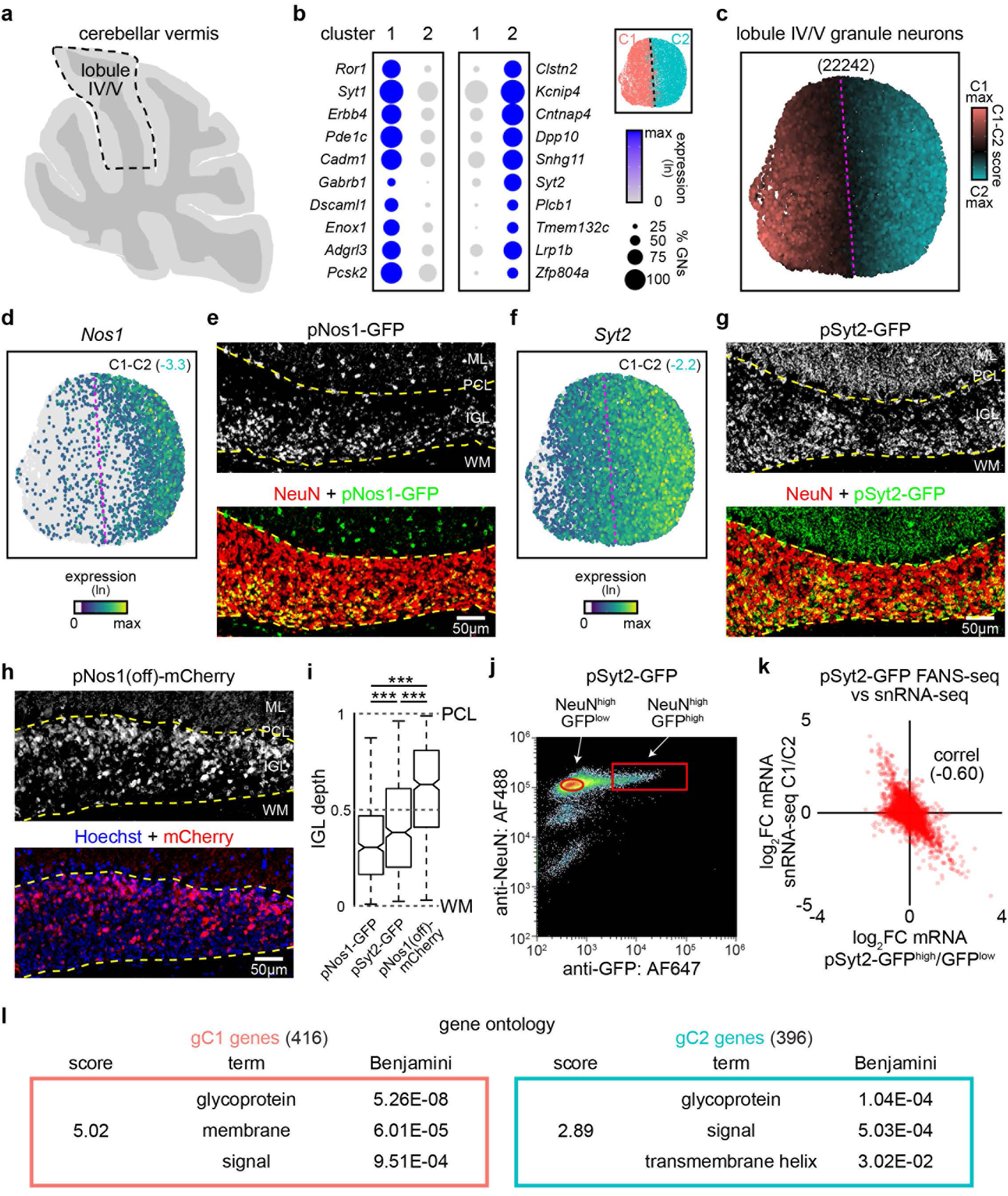
Identification of granule neuron subpopulations in cerebellar lobule IV/V. **a**, Schematic of a sagittal section of the cerebellar vermis in an adult mouse. The dashed line indicates lobule IV/V. **b**, Dot plot of selected marker genes showing the average normalized expression and percentage of granule neurons in lobule IV/V expressing each gene within granule neuron clusters^12^. Inset: UMAP plot of granule neuron clusters 1 (C1) and 2 (C2), with the dotted line denoting the separation between clusters. **c**, Relative C1 and C2 scores overlaid onto the UMAP from **b**, indicating the graded expression of genes that distinguish granule neuron subpopulations. **d**, Expression of *Nos1* in lobule IV/V granule neurons visualized with the UMAP plot in **c**, with the log fold change in expression in C1 compared with C2 neurons indicated (C1-C2). **e**, Labeling of cerebellar lobule IV/V in pNos1-GFP mice with antibodies against GFP and the granule neuron marker NeuN. Layers within the cerebellar cortex including the molecular layer (ML), Purkinje cell layer (PCL), internal granule layer (IGL), and white matter (WM) are shown, with dashed lines indicating the IGL boundary. **f**, Expression of *Syt2* in lobule IV/V granule neurons visualized with the UMAP plot in **c**, with the log fold change in expression in C1 compared with C2 neurons indicated (C1-C2). **g**, **h**, Labeling of cerebellar lobule IV/V in pSyt2-GFP mice with antibodies against GFP and NeuN (**g**) or in pNos1(off)-mCherry mice with an antibody against mCherry and Hoechst DNA dye (**h**). **i**, Localization of fluorescent protein (FP)-labeled cells in the IGL as a function of distance relative to the PCL (set at 1) and the WM (set at 0), from **e**, **g**, and **h** (\*\*\**P* < 0.001, Kruskal-Wallis test with Dunn’s post hoc test with Šidák correction, *n* = 512, 649, 557 cells in 2 mice for pNos1-GFP, pSyt2-GFP, pNos1(off)-mCherry). Box plots show median, quartiles (box), and range (whiskers). **j**, Fluorescence-activated nuclei sorting (FANS) procedure for isolating granule neurons using antibodies targeting GFP and NeuN. The fluorescence intensities of labeled nuclei and the sorting gates are shown for GFP^high^/NeuN^high^ (red rectangle) and GFP^low^/NeuN^high^ (red circle) subpopulations from pSyt2-GFP mice. **k**, Comparison of log fold changes in gene expression between GFP^high^ and GFP^low^ subpopulations in pSyt2-GFP mice with log fold changes in gene expression those between C1 and C2 clusters from snRNA-seq analyses. The correlation coefficient (correl) is indicated. **l**, Gene Ontology analyses of genetically-defined C1 (gC1) and C2 (gC2) marker genes from FANS-RNA-seq analyses, with DAVID group enrichment scores for functional annotation clusters.

To characterize these granule neuron subpopulations further, we obtained transgenic mouse lines from GENSAT that express GFP fluorescent protein under the control of the *Nos1* or *Syt2* promoter (pNos1-GFP or pSyt2-GFP)^20^, with *Nos1* and *Syt2* serving as markers enriched in C2 granule neurons relative to C1 neurons. Immunohistochemical analyses revealed that these GFP-labeled granule neurons were enriched in the inner-zone of the IGL, adjacent to the white matter (Fig. 1d-g). We also employed an intersectional strategy using mice expressing Cre recombinase under the *Nos1* promoter and FlpO recombinase under the glutamatergic neuron-specific *Vglut1* promoter. In these mice, delivery of an AAV reporter encoding Cre_off_-Flp_on_-mCherry^21^ selectively labeled the C1 granule neuron subpopulation (pNos1(off)-mCherry; Methods). These mCherry-labeled C1 neurons were enriched in the outer-zone of the IGL (Fig. 1h). Quantification of granule neuron soma locations across these three transgenic mouse lines confirmed labeling of different zones in the IGL (Fig. 1i), indicating that genetically-defined granule neuron subpopulations labeled using the *Nos1* or *Syt2* promoter are spatially distinguishable in the cerebellar cortex. Consistent with these findings, publicly available *in situ* hybridization data showed that expression of *Syt1* and *Cadm1*, markers of C1 subpopulation, was enriched in the outer-zone of the IGL near the Purkinje cell layer, whereas the expression of *Cntnap4*, a marker of C2 subpopulation, was enriched in the inner-zone of the IGL (Extended Data Fig. 1a, b)^22^.

We next profiled the transcriptomes of granule neuron subpopulations in our transgenic animals by performing fluorescence-activated nuclei sorting (FANS) using antibodies against GFP or mCherry together with the granule neuron marker NeuN, followed by RNA-seq analyses (Fig. 1j; Extended Data Fig. 1c). We found that gene expression differences between GFP^high^ and GFP^low^ granule neuron subpopulations in pNos1-GFP or pSyt2-GFP mice were inversely correlated with the gene expression differences between C1 and C2 neurons identified by snRNA-seq, with C2 and C1 markers preferentially expressed in the GFP^high^ and GFP^low^ subpopulations, respectively (Fig. 1k and Extended Data Fig. 1d, e). By contrast, this relationship was reversed in pNos1(off)-mCherry mice, with mCherry^high^ and mCherry^low^ subpopulations corresponding to the C1 and C2 expression profiles, respectively (Extended Data Fig. 1d, e). These results demonstrate that our transgenic lines reliably label the C1 and C2 granule neuron subpopulations.

Using these mouse genetic lines, we identified a common set of marker genes that were reproducibly differentially expressed across the datasets, including 416 genes enriched in the genetically-defined C1 (gC1) subpopulation and 396 genes enriched in the C2 (gC2) subpopulation (Supplementary Table 1). Gene ontology analysis of these marker genes showed that both C1 and C2 granule neuron subpopulations were enriched for genes associated with transmembrane or signaling functions, such as *Nos1*, *Syt1*, and *Syt2* (Fig. 1l).

In addition, we found that immediate-early genes (IEGs), including *Fos* and *Npas4*, which are rapidly and robustly induced in granule neurons during sensorimotor behavior or depolarization^10, 23^, exhibited lower expression in the C2 subpopulation labeled by pNos1-GFP or pSyt2-GFP compared with the C1 subpopulation labeled by pNos1(off)-mCherry (Extended Data Fig. 1f). This enrichment of IEGs in C1 compared with C2 neurons was also observed in snRNA-seq data (Extended Data Fig. 1g). These results suggest that C1 granule neurons are more active than C2 granule neurons in mice maintained under standard home cage conditions. Together, our findings indicate that the C1 and C2 granule neuron subpopulations, each enriched in separate zones of the IGL, may differentially respond to sensorimotor activity.

### The C2 gene expression program predominates during early postmitotic development

The identification of transcriptionally and spatially distinguishable granule neuron subpopulations led us to investigate when these subpopulations emerge during cerebellar development. We first used *in vivo* electroporation to label granule neuron precursors exiting the cell cycle with an mCherry-NLS expression vector and tracked their postmitotic differentiation through defined developmental stages, including dendrite growth at 4 days after electroporation, synapse formation at day 8, and synapse maturation at day 14 (Fig. 2a)^11, 24, 25^. We isolated developing granule neurons by FANS using antibodies against mCherry and NeuN, followed by RNA-seq analyses, and identified 8295 genes that were differentially expressed across these developmental stages (Supplementary Table 2). In addition, we performed FANS followed by snRNA-seq analyses to characterize postmitotic granule neurons from cerebellar lobule IV/V of P16 juvenile mice (Extended Data Fig 2a; Methods). We identified four major granule neuron clusters by unsupervised clustering and then projected the gene expression profiles of electroporated granule neurons at defined developmental stages onto the P16 UMAP (Fig. 2b, c). These analyses revealed that the major granule neuron clusters corresponded to postmitotic developmental stages (Fig. 2b, c and Extended Data Fig. 2b, c; Methods).

**Fig. 2.**
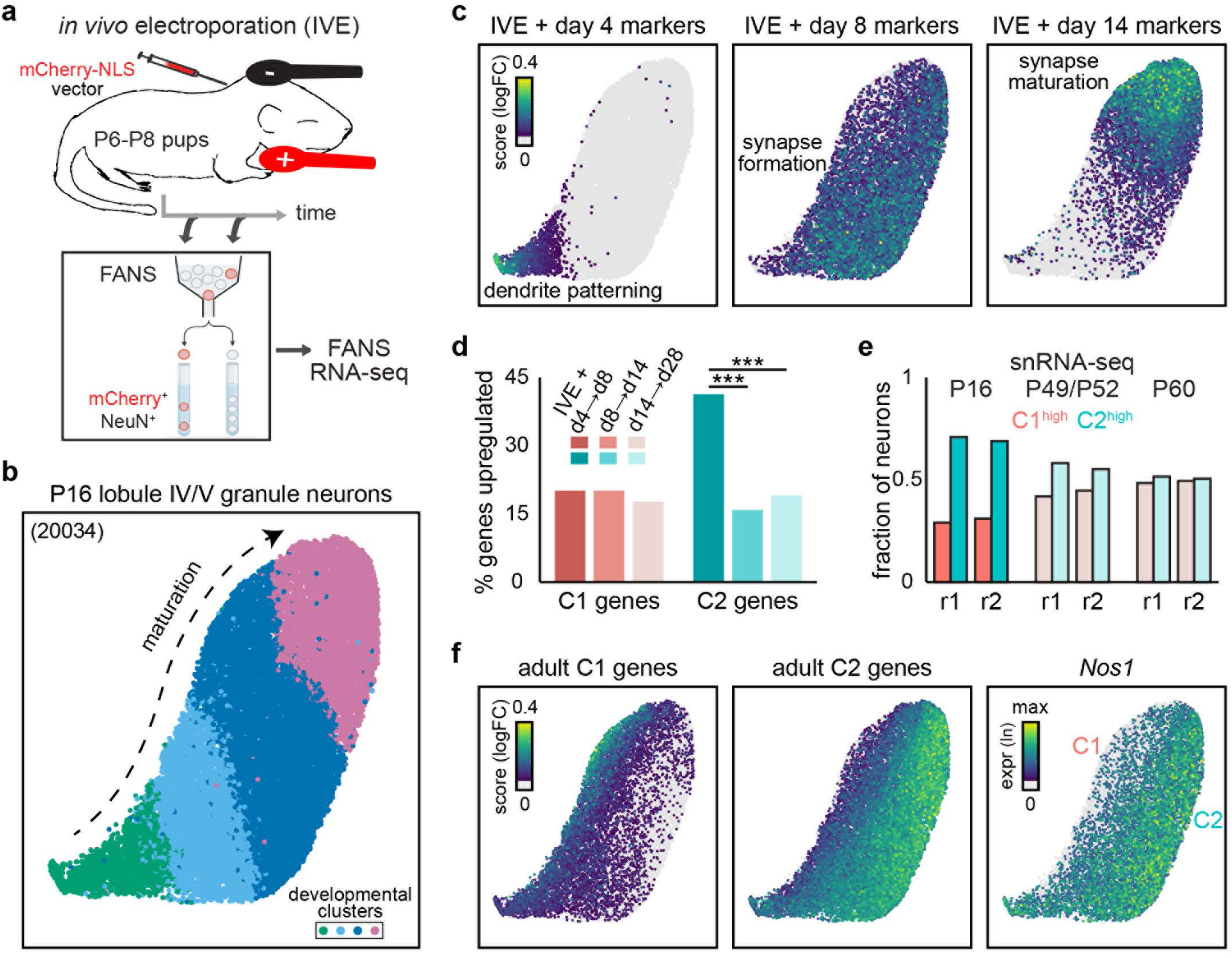
Granule neurons preferentially upregulate C2 gene expression in early development. **a**, The *in vivo* electroporation (IVE) approach used for transcriptional profiling of synchronously differentiating granule neurons at defined stages. Mouse pups were electroporated at P6-P8 with an mCherry-NLS expression plasmid and mCherry-labeled granule neuron nuclei were isolated by FANS using antibodies against mCherry and NeuN, followed by RNA-seq analyses. **b**, **c**, UMAP plot of lobule IV/V granule neuron nuclei from P16 mice (**b**). Expression scores for genes enriched at day 4, 8, or 14 after electroporation, which correspond to periods of dendrite patterning, synapse formation, and synapse maturation, respectively^11, 24^, overlaid on the UMAP plot (**c**). The granule neuron clusters identified in **b** align with stages of postmitotic differentiation. **d**, Fraction of C1 or C2 marker genes upregulated (log_2_FC > 0.585) during the indicated developmental windows after electroporation (\*\*\**FDR* < 0.001, pairwise Fisher’s exact test with Benjamini-Hochberg correction). **e**, Fraction of C1^high^ versus C2^high^ granule neurons at P16, P49/P52, and P60, as in Extended Data Fig. 2d. C2^high^ granule neurons predominated at P16, while the C1^high^ and C2^high^ granule neurons were present at comparable proportions by P60. Two biological replicates (r1 and r2) are shown. **f**, Expression scores for C1 and C2 marker genes and expression of the C2 marker *Nos1* overlaid on the UMAP plot from P16 mice.

We next asked when and how the C1 and C2 gene expression programs emerge during postmitotic differentiation by tracking the expression of C1 and C2 markers, which were defined in adulthood, from day 4 to day 28 after electroporation. Remarkably, many C2 marker genes were robustly upregulated between days 4 and 8, coinciding with the period of synapse formation, while C1 marker genes changed more gradually across development (Fig. 2c, d). These findings suggest that the C2 gene expression program is preferentially upregulated during the developmental window when granule neurons undergo synaptogenesis. We next asked whether this preferential upregulation led to a predominance of the C2 gene expression program among individual granule neurons during this developmental window. In our snRNA-seq dataset from P16 juvenile mice, we found that more than 70% of postmitotic granule neurons showed higher C2 than C1 gene expression (Fig. 2e, f and Extended Data Fig. 2d). By P60, however, the proportion of granule neurons with higher C2 gene expression had declined, resulting in comparable numbers of granule neurons with higher C1 or C2 gene expression (Fig. 2e and Extended Data Fig. 2d). Together, these results suggest that granule neurons preferentially expressing the C2 program predominated during early postmitotic development, while C1 and C2 granule neurons became more comparably represented in adulthood.

### Downregulation of the C2 program in a subset of neurons during late postnatal development

We further investigated the regulation of the C1 and C2 gene expression programs during late postnatal development, from the juvenile to the young adult stage. To compare these programs, we sought to identify a marker whose expression could stratify granule neurons by relative C1 and C2 gene expression at both stages. Notably, *Rbfox3*, which encodes the NeuN protein and is a well-established marker of granule neurons among cerebellar cell types, exhibited graded expression across the granule neuron subpopulations in both the P16 and P60 snRNA-seq datasets. Granule neurons with relatively lower *Rbfox3* expression showed higher expression of C1 genes, while those with relatively higher *Rbfox3* expression showed higher expression of C2 genes (Extended Data Fig. 3a). Given these findings, we performed FANS using an antibody against NeuN to stratify granule neurons based on NeuN protein levels. We first gated NeuN-positive granule neurons to exclude other cerebellar cell types, including glia and inhibitory neurons, and then stratified granule neurons into fractions containing the bottom, middle, and top 30% of the NeuN fluorescence intensity distribution (Fig. 3a, b; bottom-, middle-, and top-NeuN groups). Consistent with the snRNA-seq results, FANS-RNA-seq analyses showed that the bottom-NeuN group was enriched for C1 genes, while the top-NeuN group was enriched for C2 genes (Fig. 3c and Extended Data Fig. 3a).

**Fig. 3.**
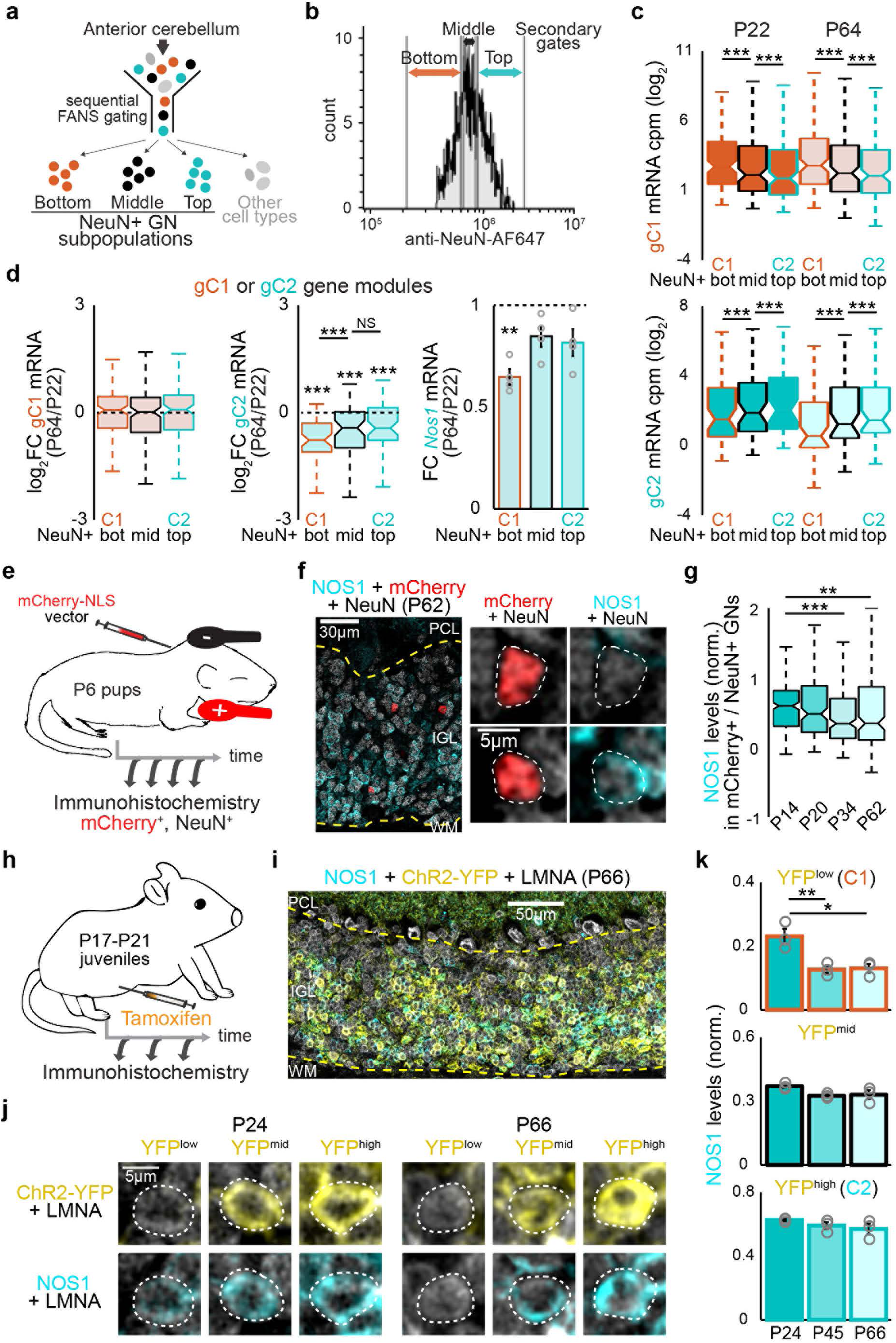
The C2 program is downregulated in a subset of granule neurons in late development. **a**, FANS procedure using a primary gate to enrich for NeuN-positive granule neurons^44^ and a secondary gate to stratify granule neuron subpopulations by relative NeuN levels. **b**, Secondary FANS gates to separate the bottom, middle, and top 30% of NeuN fluorescence intensity among NeuN-positive granule neurons. **c**, Expression of genetically-defined C1 (gC1) and C2 (gC2) marker genes in the bottom, middle, and top fractions of NeuN-positive granule neurons from the anterior cerebellum at P22 and P64 (two-sided Wilcoxon signed-rank test, *n* = 125, 54 genes for gC1, gC2). **d**, Left and middle, fold changes in C1 and C2 marker gene expression in NeuN-stratified granule neurons as in **c** from P22 to P64. C2 marker genes showed an overall decrease in expression during late postnatal development (two-sided one-sample Wilcoxon signed-rank test against 0, *n* = 54 genes), with the greatest decrease in the bottom-NeuN, C1-enriched subpopulation (two-sided Wilcoxon signed-rank test). Right, fold changes in *Nos1* expression from P22 to P64 in NeuN-stratified granule neurons (two-sided one-sample *t*-test against 0, *n* = 4 biological replicates). **e**, **f**, The IVE approach as in Fig. 2a was used to deliver an mCherry-NLS expression plasmid to P6 pups and label synchronously differentiating granule neurons (**e**). At P14, P20, P34, and P62, electroporated granule neurons were visualized using antibodies against mCherry, NeuN, and NOS1. A labeled region in cerebellar lobule IV/V at P62 is shown, with zoomed-in examples of mCherry- and NeuN-positive granule neurons shown on the right. The PCL, IGL, and WM are indicated, with dashed lines marking the IGL boundary (**f**). **g**, Distribution of NOS1 protein levels in electroporated granule neurons at the indicated postnatal ages as in Extended Data Fig. 3b (two-sided Wilcoxon rank-sum tests against the P14 condition with Benjamini-Hochberg correction, *n* = 266, 152, 211, 158 cells for P14, P20, P34, P62). **h**-**j**, Fate-tracing procedure with tamoxifen injections from P17 to P21 in juvenile transgenic mice that express ChR2-YFP in granule neurons a pNos1-CreERT2 dependent manner (Methods), followed by immunohistochemistry using antibodies against GFP/YFP, LMNA, and NOS1 at P24, P45, and P66 (**h**). Nuclear LMNA labeling delineated cell boundaries, enabling high-throughput analysis of >5000 cells per sample (Methods). A labeled region in lobule IV/V at P66 is shown, with dashed lines indicating the IGL boundary (**i**). LMNA-labeled IGL cells with increasing ChR2-YFP intensities at P24 and P66 (**j**). The dashed lines denote the cell boundaries. **k**, NOS1 protein levels in LMNA-positive IGL cells stratified by ChR2-YFP intensity, shown as in **j**, at P24, P45, and P66 (one-way ANOVA with Dunnett’s post hoc test against the P24 condition, *n* = 3 biological replicates). Box plots in **c**, **d**, left and middle, and **g** show median, quartiles (box), and range (whiskers). Data in **d**, right, and **k** show mean and error bars denote s.e.m. \**P* < 0.05, \*\**P* < 0.01, \*\*\**P* < 0.001.

We next examined how these C1 and C2 gene expression programs changed within the NeuN-stratified groups between P22, when all postmitotic granule neurons have integrated into cerebellar circuits, and P64, when mice have reached adulthood. Across the NeuN-stratified groups, the expression of C1 genes exhibited little or no overall change from P22 to P64, while C2 gene expression decreased during this period (Fig. 3d). Importantly, the reduction in C2 gene expression was greatest in the bottom-NeuN group, which is enriched for C1 granule neurons. This pattern was exemplified by the C2 marker gene *Nos1*, whose expression showed the greatest reduction over this period in the bottom-NeuN group (Fig. 3d). These findings suggest that C1 granule neuron identity may be refined during late postnatal development through downregulation of the C2 program, rather than through further changes in the C1 program.

To visualize these late developmental changes in the C2 program in the cerebellar cortex, we combined immunohistochemistry for the C2 marker NOS1 with labeling of synchronously differentiating granule neurons by *in vivo* electroporation of an mCherry-NLS expression vector at P6^11, 24, 25^. We then examined these synchronously differentiating granule neurons at P14, 8 days after electroporation when the C2 gene expression program was upregulated, and subsequently at P20, P34, and P62 (Figs. 2d and 3e). At all ages, NOS1 protein levels varied substantially among mCherry-labeled granule neurons, which comprise both C1 and C2 subpopulations (Fig. 3f and Extended Data Fig. 3b). Interestingly, an increasing fraction of mCherry-labeled granule neurons exhibited lower NOS1 protein levels over this time period, consistent with our findings of decreased *Nos1* mRNA expression (Fig. 3d, g).

To determine which granule neuron subpopulation underwent this NOS1 protein downregulation, we performed fate-tracing by inducibly labeling granule neurons with ChR2-YFP under the control of the *Nos1* promoter in juvenile mice from P17 to P21, when NOS1 expression was high across a broader fraction of granule neurons than in adulthood (Fig. 3g, h and Extended Data Fig. 3b-d; Methods). This strategy permanently labeled granule neurons with YFP based on *Nos1* promoter activity during this early time window, allowing us to assess endogenous NOS1 levels in these fate-traced cells at later stages (Fig. 3i). We thus stratified cells in the IGL by YFP fluorescence intensity into the YFP^high^, YFP^mid^, and YFP^low^ groups (Fig. 3j). NOS1 protein levels declined primarily in the YFP^low^ group between P24 and the later P45 and P66 time points (Fig. 3k). Given that lower *Nos1* promoter activity in the YFP^low^ group enriches for C1 granule neurons (Extended Data Fig. 1d, e), these findings support our observations that C1 granule neuron identity is refined during late postnatal development through downregulation of C2 markers such as NOS1.

### *In vivo* CRISPR mini-screen identifies genetic regulators of granule neuron subpopulations

We next investigated the mechanisms underlying these gene expression changes at defined developmental stages by performing genome-wide profiling of gene regulatory enhancer activity. Using FANS-ChIP-seq analyses with an antibody against H3K27ac, a histone mark deposited at active enhancers, we identified enhancers that were differentially regulated across development (Fig. 4a, b). Interestingly, DNA binding motifs for the transcription factors ZEB1/2, TEAD1, ETV1, ZIC2, and RORA/C were selectively enriched at enhancers upregulated between day 4 and day 8 after electroporation, whereas motifs for IEGs including FOS and NR4A1 were enriched between day 8 and day 14 (Fig. 4c). These results suggest that C1 and C2 granule neurons may be regulated by specific transcription factors over different developmental windows.

**Fig. 4.**
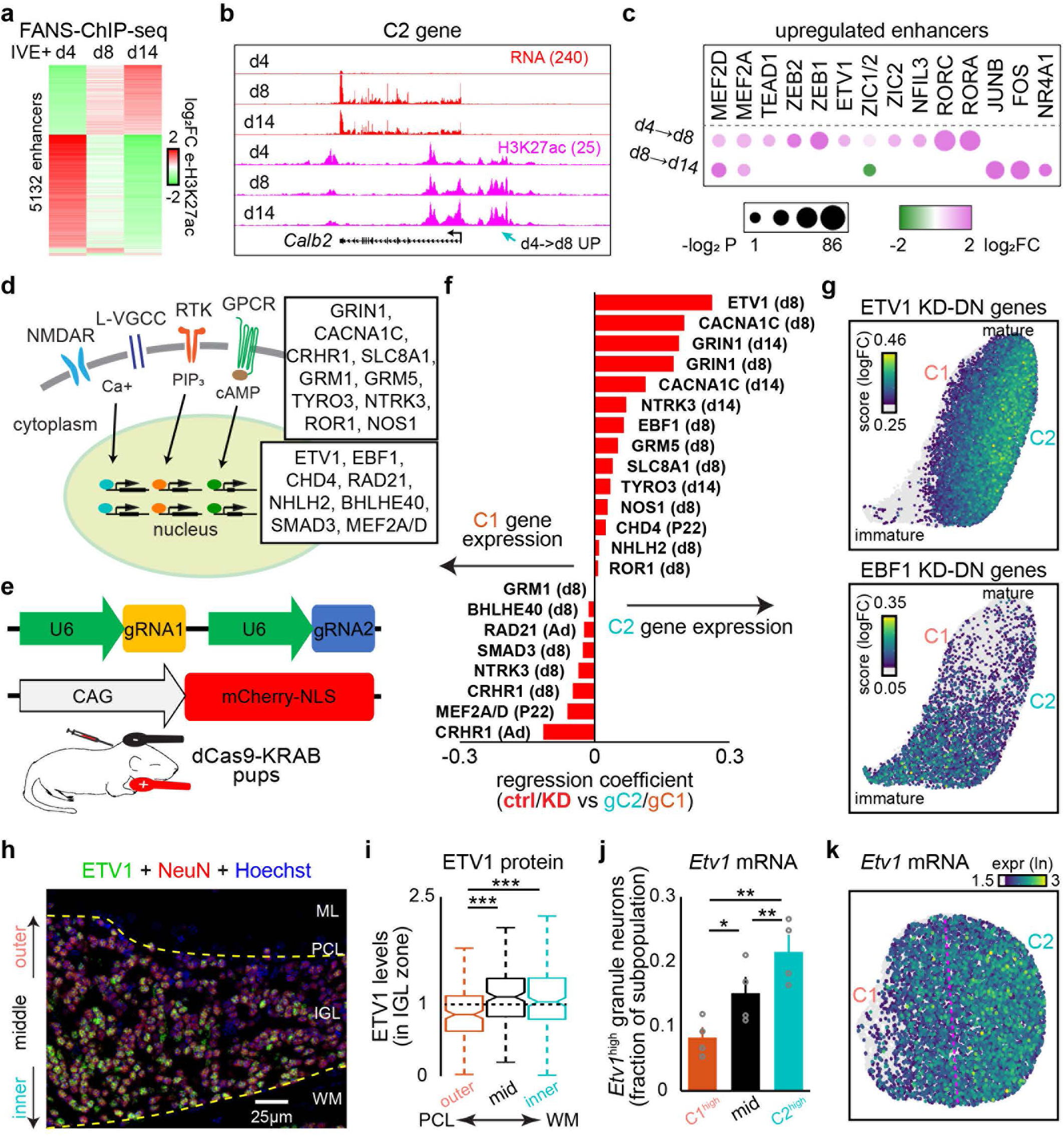
*In vivo* CRISPR mini-screen identifies regulators of granule neuron subpopulations. **a**, Log_2_ fold changes in H3K27ac levels at significantly differentially active regulatory enhancers (FDR < 0.05, two-sided *P* value from a likelihood-ratio test using a negative binomial generalized linear model with Benjamini-Hochberg correction, *n* = 2, 3, 3 biological replicates) in granule neurons at day 4, 8, or 14 after electroporation. Granule neurons were isolated using FANS and subjected to ChIP-seq analyses using an H3K27ac antibody. Data are mean centered. **b**, UCSC genome browser tracks showing gene expression or H3K27ac levels at the *Calb2* locus, a C2 marker gene, in granule neurons at day 4, 8, or 14 after electroporation. The cyan arrow denotes an upstream enhancer with increased H3K27ac levels from day 4 to day 8. **c**, Transcription factor binding motifs enriched at enhancers with increased H3K27ac levels during specific stages of granule neuron development. **d**, Schematic of transmembrane receptors that respond to extracellular cues and trigger intracellular signaling, which may activate transcriptional regulators and gene expression programs in the nucleus. Candidate regulators tested in the *in vivo* mini-screen, including these receptors and nuclear factors, are indicated. **e**, Design of the plasmid DNA used for *in vivo* electroporation to label granule neurons and induce gene silencing. **f**, Regression coefficients showing the effects of knockdown or knockout of 19 candidate regulators on the gene expression programs of genetically-defined C1 and C2 granule neuron subpopulations, including 14 perturbations generated in this study and others from previous datasets^26–29^, relative to the gene expression differences between these subpopulations in pSyt2-GFP mice. **g**, Expression scores of genes downregulated upon ETV1 knockdown (top) or EBF1 knockdown (bottom) compared to control neurons, overlaid onto P16 UMAP plot. **h**, Image of the cerebellar cortex from adult mice labeled with the ETV1 (green) and NeuN (red) antibodies together with the Hoechst DNA dye (blue). Layers of the cerebellar cortex with dashed lines indicating the IGL boundaries. **i**, ETV1 protein levels in the outer, middle, and inner thirds of the IGL, as shown in **h**. ETV1 protein was enriched in the inner two-thirds of the IGL (Kruskal-Wallis test with Dunn’s post hoc test with Šidák correction, *n* = 183, 194, 212 granule neurons in outer, middle, inner from 2 mice). The dashed line indicates the median level across the granule neuron population. **j**, Fraction of granule neuron in the top, middle, or bottom C1-C2 score groups from P49, P52, and P60 mice that express high levels of *Etv1* mRNA. C2 granule neurons expressed higher *Etv1* mRNA levels than the other subpopulations (repeated-measures one-way ANOVA with Tukey-Kramer post hoc test, *n* = 4 biological replicates). **k**, *Etv1* mRNA expression visualized on a UMAP plot of cerebellar granule neurons from P60 mice as in Fig. 1c. Box plots in **i** show median, quartiles (box), and range (whiskers). Data in **j** show mean and error bars denote s.e.m. \**P* < 0.05, \*\**P* < 0.01, \*\*\**P* < 0.001.

To identify factors that functionally regulate the gene expression programs associated with C1 and C2 granule neurons, we performed an *in vivo* CRISPR mini-screen. We examined the functions of 19 candidate regulators including transcriptional and chromatin regulators that were enriched in the C1 or C2 subpopulations, expressed during specific stages of granule neuron development such as the period of synaptogenesis, or curated from our previous datasets^26–29^, including ETV1, EBF1, CHD4, RAD21, NHLH2, BHLHE40, SMAD3, and MEF2A/D (Fig. 4d). We also tested signaling factors enriched in granule neurons that may couple extrinsic cues to calcium and other intracellular pathways to regulate gene transcription, including GRIN1, CACNA1C, CRHR1, SLC8A1, GRM1/5, TYRO3, NTRK3, ROR1, and NOS1 (Fig. 4d). To induce knockdown of these targets *in vivo*, we electroporated transgenic mouse pups expressing dCas9-KRAB with an mCherry expression plasmid together with a vector encoding sgRNAs targeting gene promoters or control sequences (Fig. 4e)^29, 30^. We then isolated NeuN and mCherry-labeled granule neurons using FANS and performed RNA-seq at day 8 or day 14 after electroporation. The knockdown efficiency of each sgRNA was validated, and target genes with consistent depletion were subsequently subjected to transcriptome-wide analyses (Extended Data Fig. 4a).

We assessed how the knockdown of each mini-screen target affected the C1/C2-associated gene expression programs. We identified several robust regulators of the C2 program, including the ETS variant transcription factor 1 (ETV1), the glutamate ionotropic receptor NMDA type subunit 1 (GRIN1), and the L-type voltage-gated calcium channel subunit α-1C (CACNA1C) (Fig. 4f). Notably, ETV1 knockdown reduced the expression of the largest number of C2-associated genes compared with other mini-screen targets, while having limited effects on the C1 program (Extended Data Fig. 4b-d). Consistent with this selective effect, ETV1-dependent genes were enriched in C2 granule neurons in snRNA-seq analyses of both P16 and adult mice (Fig. 4g, top and Extended Data Fig. 4e). By contrast, other transcription factors tested in the mini-screen had modest or minimal effects on the C2 program. For example, EBF1-dependent genes were selectively expressed in immature postmitotic granule neurons and were downregulated during development (Fig. 4g, bottom). Among mini-screen targets involved in signaling, GRIN1 regulated a subset of genes associated with C2 neurons, while CRHR1 regulated a group of genes selective for C1 granule neurons (Fig. 4f and Extended Data Fig. 4c, d, f, g). Together, these analyses identified potential signaling and transcriptional regulators of granule neuron subpopulations in the cerebellum.

Given that ETV1 plays a prominent and specific role in promoting the transcriptional program of C2 granule neurons, we next examined whether ETV1 is differentially expressed between C1 and C2 granule neurons. Immunohistochemical analyses using an antibody against ETV1 revealed that ETV1 protein was enriched in granule neurons located in the middle and inner thirds of the IGL, where C2 granule neurons are preferentially localized (Figs. 1i and 4h, i). Consistently, *Etv1* mRNA was also enriched in C2 granule neurons and depleted in C1 granule neurons (Fig. 4j, k). These findings suggest that the enrichment of ETV1 in C2 granule neurons may contribute to the establishment of their transcriptional identity.

We also examined the contributions of the transmembrane proteins GRIN1 and CACNA1C, which regulate a smaller subset of genes compared to ETV1, in the specification of C2 granule neurons (Fig. 4f and Extended Data Fig. 4c, d). Because GRIN1 and CACNA1C are subunits of calcium ion channels, we reasoned that they might act upstream of ETV1 to promote gene transcription in the nucleus. Analysis of C2 marker genes such as *Syt2*, *Kcnip4*, and *Plekhg3* revealed that combined knockdown of GRIN1 and CACNA1C reduced their expression, similar to the effect of ETV1 knockdown (Extended Data Fig. 4h-j). Interestingly, combined knockdown of GRIN1 and CACNA1C led to greater reductions in *Syt2*, *Kcnip4*, and *Plekhg3* expression than either knockdown alone (Extended Data Fig. 4h-j). These effects extended genome-wide, with ETV1-dependent genes more strongly downregulated by the combined knockdown of GRIN1 and CACNA1C (Extended Data Fig. 4k), suggesting that these ion channels act through parallel calcium signaling pathways to regulate ETV1-dependent gene expression. Together, these results indicate that the transcriptional identity of C2 granule neurons is driven by ETV1 and by NMDARs and L-type VGCC signaling during development.

### Calcium signaling and ETV1 activates regulatory enhancers associated with C2 granule neurons

Having identified the transcription factor ETV1 as a regulator of the C2 gene expression program, we examined genomic regions associated with C1 and C2 marker genes to determine whether ETV1 serves as the primary factor in the regulation of these genes. We first analyzed published <u>s</u>ingle <u>n</u>uclei ATAC-seq (snATAC-seq) datasets from anterior cerebellar granule neurons in adult mice^31, 32^ and found that chromatin accessibility patterns at C1 and C2 marker genes distinguished these two subpopulations on the UMAP projection of snATAC-seq data (Fig. 5a, left; Methods).

**Fig. 5.**
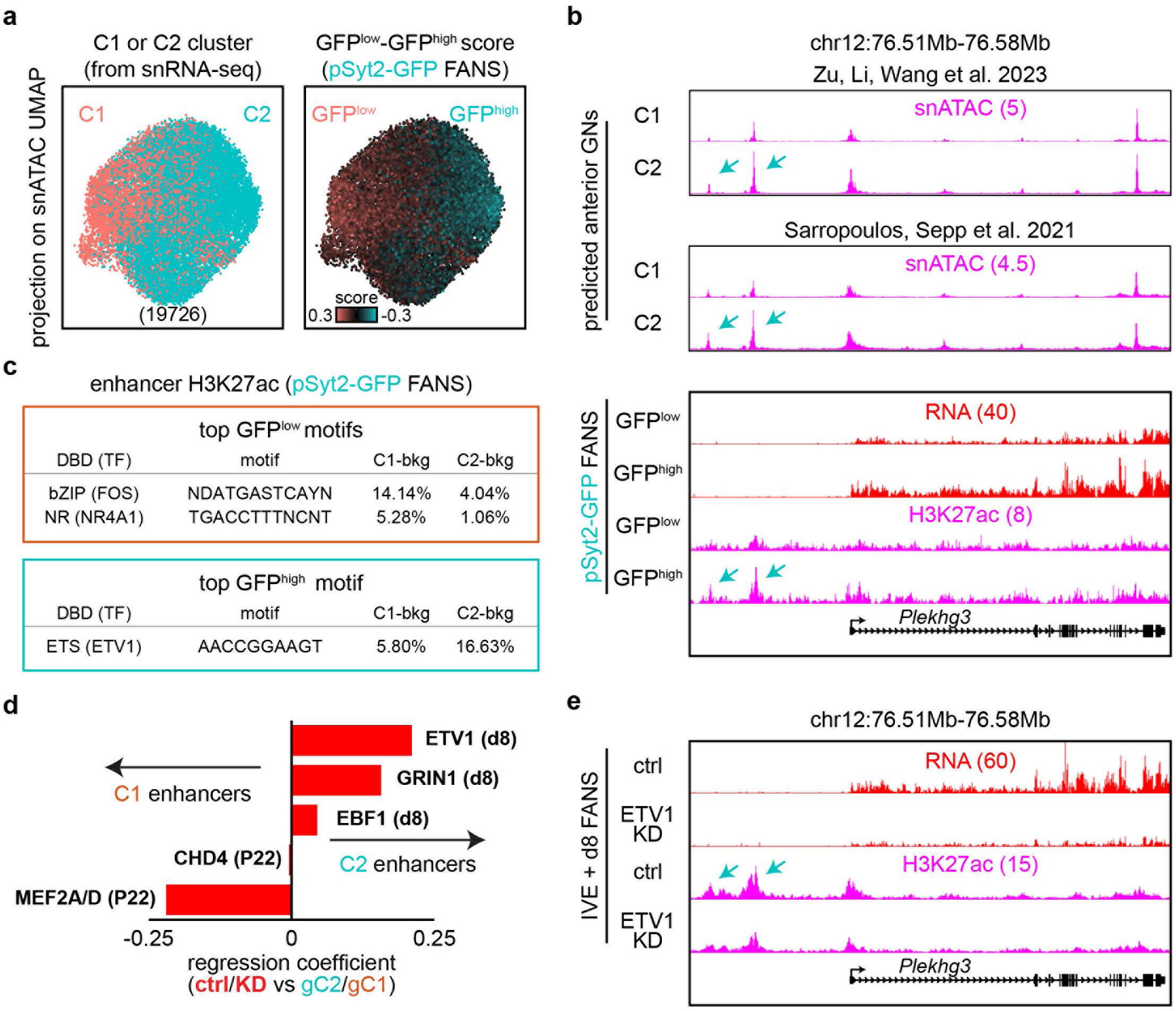
ETV1 is a major regulator of active enhancers in C2 granule neurons. **a**, C1 and C2 annotations derived from integrative snRNA-seq analyses (left) or GFP^low^-GFP^high^ module scores from pSyt2-GFP mice (right) overlaid onto the snATAC-seq UMAP plot of anterior granule neurons in adult cerebellum^31^. **b**, UCSC genome browser tracks showing pseudobulk chromatin accessibility from published datasets^31, 32^ in predicted C1 and C2 granule neurons (top), and gene expression and H3K27ac levels in GFP^low^ and GFP^high^ granule neurons isolated from the anterior cerebellar lobules of pSyt2-GFP mice (bottom) at the *Plekhg3* locus, a C2 marker gene. **c**, Transcription factor binding motifs enriched in enhancers selectively active in GFP^low^ (C1) or GFP^high^ (C2) granule neurons in adult mice (FDR < 0.1, two-sided *P* value from a likelihood-ratio test using a negative binomial generalized linear model with Benjamini-Hochberg correction, *n* = 4 biological replicates). **d**, Regression coefficients showing the effects of knockdown or knockout of candidate regulators on enhancers associated with genetically-defined C1 and C2 granule neuron subpopulations, relative to the differences in enhancer activity between these subpopulations in pSyt2-GFP mice. **e**, UCSC genome browser tracks showing gene expression and H3K27ac levels at the *Plekhg3* locus, as in **b**, in Etv1 knockdown or control granule neurons at day 8 after electroporation. Cyan arrows in **b**, **e** denote upstream enhancers.

We therefore hypothesized that genome-wide differences in chromatin accessibility reflect differences in the activity of regulatory enhancers between C1 and C2 granule neurons. To test this, we performed H3K27ac ChIP-seq analyses on GFP^low^ and GFP^high^ granule neurons isolated from the anterior cerebellum of pSyt2-GFP mice, which correspond to the C1 and C2 subpopulations, respectively (Fig. 1). When the accessibility profiles of enhancers with differential H3K27ac levels were projected onto the snATAC-seq UMAP plot, enhancers with higher H3K27ac levels in GFP^low^ and GFP^high^ granule neurons corresponded to the C1 and C2 subpopulations, respectively (Fig. 5a, right). For example, at the locus of the C2 marker gene *Plekhg3*, upstream regulatory enhancers showed greater chromatin accessibility and higher H3K27ac levels in GFP^high^ C2 granule neurons than in GFP^low^ C1 neurons (Fig. 5b). More broadly, enhancers near C1-enriched genes showed higher H3K27ac levels in GFP^low^ C1 granule neurons, while enhancers near C2-enriched genes showed higher H3K27ac levels in GFP^high^ C2 granule neurons (Extended Data Fig. 5a, b). These results suggest that distinct sets of active regulatory enhancers define the C1 and C2 transcriptional programs in the cerebellum.

Strikingly, DNA motif analyses of C2 regulatory enhancers from both ChIP-seq and snATAC-seq datasets revealed that the ETS consensus motif was the top enriched sequence (Fig. 5c and Extended Data Fig. 5c). These findings are consistent with our genetic mini-screen showing that ETV1 acts as a major regulator of the transcriptional identity of C2 granule neurons (Fig. 4). In contrast, we observed that bZIP and NR motifs were enriched at C1 enhancers, consistent with higher mRNA expression of *Fos/Fosl2* and *Nr4a1/2/3* in C1 granule neurons (Fig. 5c and Extended Data Figs. 1f, g and 5b). These findings suggest that distinct sets of transcription factors may act through their target enhancers to regulate the gene expression programs defining granule neuron subpopulations, with C2 enhancers notably enriched for the ETV1 consensus motif.

We next determined whether ETV1 functionally regulates C2 enhancers during development by analyzing our recent H3K27ac ChIP-seq datasets from ETV1 knockdown granule neurons^29^. We found that ETV1 robustly promoted the activity of C2 granule neuron enhancers (Fig. 5d and Extended Data Fig. 5d). Indeed, at the locus of the C2 marker gene *Plekhg3*, ETV1 knockdown reduced the activity of upstream enhancers as well as the levels of *Plekhg3* mRNA (Fig. 5e). We also performed H3K27ac ChIP-seq analyses following the knockout of other factors identified in our genetic mini-screen including GRIN1 and EBF1, to assess their roles on enhancer activity in granule neurons. Similar to ETV1, GRIN1 knockdown selectively reduced the activity of C2 enhancers (Fig. 5d), suggesting NMDAR signaling and ETV1 together control the regulatory enhancers associated with the C2 transcriptional program. In other experiments, we also found that MEF2 transcription factors contribute to the activation of C1 enhancers (Figs. 4f and 5d).

Given the genetic link between the transcription factor ETV1 and the calcium channels GRIN1 and CACNA1C in C2 granule neurons, we hypothesized that calcium signaling through these channels may regulate ETV1 expression. To test this, we performed immunohistochemical analyses of granule neurons subjected to GRIN1 knockdown, CACNA1C knockdown, combined GRIN1 and CACNA1C knockdown, or the control condition using *in vivo* electroporation and assessed the effects on ETV1 protein levels (Extended Data Fig. 5e, f). Interestingly, knockdown of GRIN1 and CACNA1C reduced ETV1 protein levels (Extended Data Fig. 5f). However, only modest changes in *Etv1* mRNA levels were observed with GRIN1 and CACNA1C depletion (Extended Data Fig. 5g). Together, these findings suggest that the calcium channels GRIN1 and CACNA1C maintain ETV1 protein levels in granule neurons through post-transcriptional mechanisms, which may contribute to their roles in establishing C2 granule neuron identity.

### Granule neuron subpopulations are differentially activated by sensorimotor activity

The characterization of transcriptionally distinct C1 and C2 granule neuron subpopulations, their establishment during development, and their persistence into adulthood led us to investigate their functions in cerebellar control of motor behavior. We first assessed how different sensorimotor experiences engage granule neuron subpopulations in the ADCV of the cerebellum. We developed a sensitive neuronal activity reporter consisting of a short half-life GFP driven by regulatory elements for the *Npas4* gene (Methods). Mice in the home cage condition exhibited sparse labeling, with GFP-positive neurons enriched in the outer two-thirds of the IGL (Extended Data Fig. 6a-d). In contrast, mice subjected to the open field or inclined treadmill paradigm showed increased numbers of GFP-positive neurons distributed throughout the IGL (Extended Data Fig. 6a-d). We obtained similar results using FANS-RNA-seq analyses of C1 and C2 granule neurons in pNos1(off)-mCherry mice (Fig. 1h, i), in which C1 granule neurons exhibited higher *Fos* and *Npas4* expression in home cage mice, while robust expression of these genes was observed in both C1 and C2 granule neurons after the inclined treadmill paradigm (Extended Data Fig. 6e). These results suggest that granule neuron subpopulations are differentially activated across sensorimotor experiences, with C2 granule neurons being primarily activated during locomotor activity.

To determine the functions of the C2 granule neuron subpopulation, we expressed channelrhodopsin specifically in C2 granule neurons in transgenic mice (GN^C2^-ChR2-YFP) for optogenetic interrogation, using the inducible labeling strategy described above but with labeling initiated in adulthood (Fig. 6a, b and Extended Data Fig. 3c). We compared GN^C2^-ChR2-YFP mice with animals expressing channelrhodopsin in all cerebellar granule neurons (GN-ChR2-YFP), which we have previously characterized^10^ (Extended Data Fig. 7a). Optogenetic stimulation (optostimulation) of ADCV with a 450 nm laser activated granule neurons in this region, inducing *Fos* and *Npas4* expression in both transgenic mouse lines^10^ (Fig. 6c). Notably, optostimulation of C2 granule neurons induced *Fos* and *Npas4* expression to ∼20% of the levels observed when all granule neurons were optostimulated, consistent with the relative numbers of channelrhodopsin-expressing granule neurons in the two lines (Extended Data Fig. 7a-d).

**Fig. 6.**
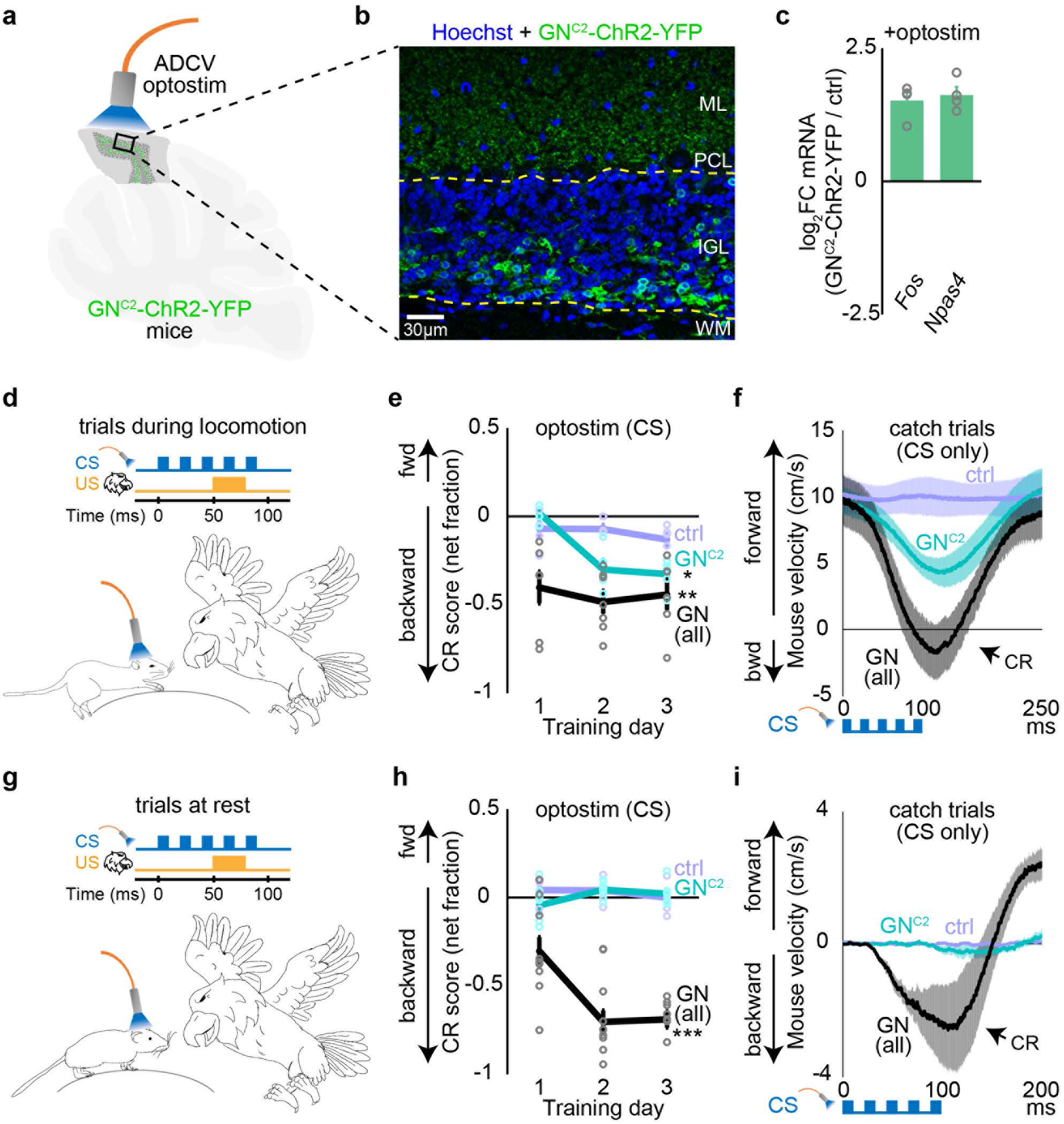
C2 granule neurons induce sensorimotor learning during locomotion. **a**, Schematic of C2 granule neurons labeled with channelrhodopsin-2 (ChR2) in adult GN^C2^-ChR2-YFP mice for optogenetic experiments. **b**, Image of the anterior dorsal cerebellar vermis (ADCV) in adult GN^C2^-ChR2-YFP mice labeled with a GFP/YFP antibody (green) and Hoechst DNA dye (blue). ChR2-YFP expression was enriched in inner-zone granule neurons. **c**, Log_2_ fold change in *Fos* and *Npas4* expression following optostimulation in GN^C2^-ChR2-YFP mice compared with control mice (two-sided one-sample *t*-test against 0, *n* = 4 biological replicates). **d**, **g**, Schematic of the delay tactile startle conditioning paradigm using head-fixed mice that were either locomoting (d) or at rest (g). Mice expressing ChR2-YFP in granule neurons, were implanted with fiber-optic cannulas over the ADCV, and optogenetically stimulated using a 450 nm laser as the conditioned stimulus (CS) paired with a tactile stimulus to the nose delivered by an eagle toy as the unconditioned stimulus (US). **e**, **f**, **h**, **i**, Control (ctrl), GN^C2^-ChR2-YFP (GN^C2^), and GN-ChR2-YFP (GN) mice, the latter expressing ChR2-YFP in all granule neurons^10^, were subjected to three days of DTSC with optostimulation as the CS. Conditioned response (CR) scores, calculated as net forward versus backward movement relative to the baseline period, are shown during locomotion (**e**) or rest trials (**h**), together with CR traces from catch trials after the third day of conditioning during locomotion (**f**) or while at rest (**i**). Mice presented a backward CR in response to optostimulation of C2 granule neurons during locomotion but not while at rest (one-way ANOVA with Dunnett’s post hoc test against the control, *n* = 5, 7, 10 mice for control, GN^C2^-ChR2-YFP, and GN-ChR2-YFP). Data in **c**, **e**, **f**, **h**, and **i** show mean and shading or error bars denote s.e.m. \**P* < 0.05, \*\**P* < 0.01, \*\*\**P* < 0.001.

To examine the roles of granule neuron subpopulations in behavior, we employed our recently developed associative learning paradigm called delay tactile startle conditioning (DTSC; Fig. 6d), which requires the ADCV^10^ and tests whether mice can associate an initially neutral sensory cue with an aversive stimulus. In this paradigm, head-fixed mice locomoting freely on a treadmill receive an unconditioned tactile stimulus (US) to the nose delivered with a motorized toy, which elicits an unconditioned startle response (Extended Data Fig. 7e). An LED cue serving as the conditioned stimulus (CS) is repeatedly paired with the tactile US, and over multiple days of training, mice acquire a conditioned backward response (CR) to the LED cue alone, indicating successful associative learning (Extended Data Fig. 7e-g)^10^.

In a variation of this paradigm, optostimulation of ADCV granule neurons replaced the LED cue as the CS and induced rapid associative sensorimotor learning, with mice acquiring a conditioned response within a few days^10^. We therefore tested whether selective activation of C2 granule neurons could induce associative learning in DTSC. Because C2 granule neurons are robustly activated by locomotion (Extended Data Fig. 6), we stratified trials based on whether animals were locomoting or at rest during CS presentation to determine whether the contribution of C2 granule neurons to associative learning depended on locomotor activity (Methods). After training, mice exhibited conditioned responses to optostimulation of C2 granule neurons during locomotion (Fig. 6d-f). In contrast, mice did not exhibit conditioned responses to C2 granule neuron optostimulation when at rest (Fig. 6g-i). However, when all ADCV granule neurons were optostimulated, trained mice exhibited conditioned responses during both locomotion and rest trials, indicating that granule neurons can induce associative learning across both behavioral conditions (Fig. 6d-i). Together, our results indicate that C2 granule neurons can induce associative learning during locomotion but not at rest and suggest that the expression of conditioned responses while mice are at rest may require broader granule neuron activity or engagement of other granule neuron subpopulations.

## Discussion

Our study defines molecular mechanisms that orchestrate gene expression programs in postmitotic neurons to diversify their cellular identities. Using a genetic mini-screen combined with RNA-seq and ChIP-seq analyses, we identify a granule neuron subpopulation enriched in the inner-zone of the internal granule layer (IGL), which we term C2 neurons, that is specified by the transcription factor ETV1. While previous studies have shown that ETV1 regulates granule neuron maturation *in vitro*^33^, our findings reveal that ETV1 selectively promotes the transcriptional program of C2 granule neurons rather than broadly inducing maturation across all granule neuron subpopulations. Moreover, calcium signaling through NMDARs and L-type VGCCs enhances ETV1 protein levels and promotes ETV1-dependent regulation of the C2 gene expression program. Calcium signaling may modulate ETV1 function either directly via post-translational modifications or through downstream effects, including changes in neuronal excitability^33, 34^. Because NMDARs and L-type VGCCs are broadly expressed across granule neuron subpopulations^22^, their selective effects on ETV1 in C2 granule neurons may arise from differences in channel activity or differences in the activation of downstream signaling pathways. The generation of postmitotic granule neurons peaks between postnatal day 5 (P5) and P10^35^, with few granule cell precursors remaining by P16. Our snRNA-seq analyses showed that at P16, ∼70% of postmitotic granule neurons preferentially expressed the C2 gene expression program, whereas granule neurons with C1 or C2 transcriptional identities were present in similar proportions by P60. Our findings indicate that this developmental shift occurs as a subset of postmitotic granule neurons downregulates the C2 gene expression program from the juvenile to the adult stage, thereby increasing the relative proportion of granule neurons with a C1 transcriptional identity by adulthood. The mechanisms underlying downregulation of the C2 program remain unclear but may involve epigenetic regulation through changes in DNA methylation, histone modifications, or chromatin organization^10, 11, 29, 36^. Interestingly, we also found that corticotropin-releasing hormone receptor 1 (CRHR1) promotes the C1 gene expression program, suggesting that GPCR-driven cAMP signaling^37^ may contribute to the maintenance of C1 transcriptional identity in adulthood.

Functionally, activation of C2 granule neurons induced associative learning selectively when mice were actively locomoting, suggesting that C2 granule neurons may contribute to learning when they are engaged by ongoing locomotor activity. Consistent with this, we found that C2 granule neurons exhibited robust activity-dependent transcriptional responses during locomotion in the treadmill paradigm, which also induces widespread cerebellar activity^38, 39^, but showed significantly lower activity-dependent transcriptional responses than C1 neurons under less active behavioral conditions. Notably, whole-cell recordings from lobule IV/V have shown that granule neurons in the inner-zone, where C2 granule neurons are enriched, have lower intrinsic excitability and are less responsive to low-frequency mossy fiber activity than those in the outer-zone, which is enriched for C1 granule neurons^40^. Together, these findings suggest that granule neuron subpopulations are differentially recruited during behavior, potentially shaping how sensorimotor information is encoded at the cerebellar input layer. While *in vivo* calcium imaging has provided the first insights into the responses of large populations of granule neurons to sensorimotor information^10, 39, 41–43^, further *in vivo* electrophysiological recordings and advances in deep imaging approaches will be needed to resolve C1 and C2 subpopulation dynamics across different depths of the cerebellar cortex during behavior.

## Acknowledgments

We thank members of the Yang Lab for helpful discussions, the Biological Imaging Facility (RRID:SCR_017767) for imaging, and the Single Cell Genomics Facility (RRID:SCR_026652). This project was supported by NIH R01NS123285 (Y.Y.) and 4DN grants U01DA053691 (T.Y.), and grants to the NSF (DMS-2235451) and Simons Foundation (MPS-NITMB-00005320) to the NSF-Simons National Institute for Theory and Mathematics in Biology (NITMB).

## Author contributions

P.V., T.Y., and Y.Y. designed the study and wrote the manuscript. P.V., S.F., and T.Y. performed RNA-seq, ChIP-seq, and FANS. P.V., and S.F. performed immunohistochemistry, *in vivo* electroporation, and behavioral analyses. Y.Y. performed bioinformatics analyses.

## Author Information

The authors declare no conflicts of interest.

## Methods

### Animals

pMath1-Cre (#011104), Cre_on_/Flp_off_-ChR2(H134R)-YFP (Ai32, #024109), dCas9-KRAB (#030000), C57BL/6J (#000664), B6D2F1/J mice (#100006), Vglut1-IRES2-FlpO-WPRE-neo (pVglut1-FlpO, #034422), pNos1-Cre (#017526), pNos1-CreERT2 (#014541) and Slc32a1-2A-FlpO-D (pSlc32a1-FlpO, #029591) mice were purchased from Jackson Laboratory, Tg(Nos1-EGFP)DP185Gsat/Mmucd (pNos1-GFP) and Tg(Syt2-EGFP)MT17Gsat/Mmucd (pSyt2-GFP) were purchased from MMRRC (Mutant Mouse Resource & Research Centers) and maintained under pathogen-free conditions. Homozygous dCas9-KRAB mice were bred with B6D2F1/J mice for *in vivo* electroporation. pVglut1-FlpO mice were bred with pNos1-Cre mice to generate pNos1-Cre/pVglut1-FlpO mice. Cre_on_/Flp_off_-ChR2(H134R)-YFP mice were bred with pMath1-Cre driver mice to generate GN-ChR2-YFP mice. Cre_on_/Flp_off_-ChR2(H134R)-YFP mice were bred with pNos1-CreERT2 and pSlc32a1-FlpO mice to generate GN^C2^-ChR2-YFP mice, in which FlpO prevents labeling of inhibitory VGAT-positive neurons. Cre recombinase activity was induced by intraperitoneal administration of tamoxifen (Sigma T-564875, 100 mg/kg for adult mice and 45mg/kg for juvenile mice; dissolved in corn oil) once daily for 5 consecutive days. Injections were administered at P17-P21 or 7-12 weeks of age, and the experiments were conducted 1 to 11 weeks after the first injection. Cre-negative littermates were used as controls for comparison with GN-ChR2-YFP and GN^C2^-ChR2-YFP mice. Both male and female mice were used and allocated to sex-matched, littermate-matched experimental groups. All animal experiments were done according to protocols approved by the Institutional Animal Care and Use Committee of Northwestern University (protocols #IS00017452 and #IS00015999) in accordance with the National Institutes of Health guidelines. Details of the animals used for experiments are listed in Supplementary Table 3.

### Antibodies

Antibodies to GFP/YFP (Abcam ab13970; HtzGFP-19F7 and HtzGFP-19C8 Memorial Sloan Kettering Cancer Center^45^), NeuN (Abcam ab177487, ab279296; Millipore MAB377), mCherry (Abcam ab205402), ETV1 (Thermo Fisher PA5-77975), LMNA (Millipore 4777), NOS1 (Abcam, ab323183), H3K27ac (Abcam ab4729), and beta III Tubulin (Abcam ab78078) for immunohistochemistry, FANS, or ChIP-seq experiments were purchased.

### Plasmid DNA

The MLM3636 vector (addgene #43860) was modified with A-U flip and stem extension to increase the stability of sgRNA as described^46^. Two sgRNAs were cloned into the modified MLM3636 vector using primers containing the target sequence listed in Supplementary Table 4. mCherry-NLS (addgene #58476) was subcloned into the pCAG vector to generate pCAG-mCherry-NLS. For the pNpas4-shGFP reporter, enhancers located at −12k (chr19:5001657-5002434) and −2k (chr19:4992897-4993291) together with the *Npas4* promoter (chr19:4989972-4990131), were inserted into the pCAG-GFP vector digested with SpeI and EcoRI. A short half-life GFP (shGFP) was generated by fusing GFP with the C-terminal 40 amino acids of mouse ODC1^47^.

### Viral vectors

The viral vector pAAV-nEF-Coff/Fon-ChR2-mCherry (Addgene #137144) using the INTRSECT viral strategy^21^ was packaged into an adeno-associated virus serotype 1 (AAV1) vector (Penn Vector Core) and delivered into pNos1-Cre/pVglut1-FlpO mice, allowing Cre and Flp-dependent intersectional expression to selectively label NOS1-negative, VGLUT1-positive granule neurons with mCherry. These animals are referred to as pNos1(off)-mCherry mice.

### Statistics

Statistical analyses were performed using Matlab, Microsoft Excel, or R. Normality of data was assessed using the Shapiro-Wilk test. For experiments in which only one or two groups were analyzed, the *t*-test was used for normal distributions. For two groups with non-normal distributions, the Wilcoxon rank-sum test (unpaired data) or Wilcoxon signed-rank test (paired data) was used. Comparisons of all samples across multiple groups were performed using analysis of variance (ANOVA) followed by Tukey’s post hoc testing (unpaired data) or repeated-measures one-way ANOVA with Tukey-Kramer post hoc test (paired data) for normal distributions, or the Kruskal-Wallis test followed by Dunn’s post hoc testing with Šidák correction for non-normal distributions. Comparisons across multiple groups to a specific condition were performed using one-way ANOVA with Dunnett’s post hoc test for normal distributions, or Wilcoxon rank-sum tests with Benjamini-Hochberg correction (unpaired data) or Wilcoxon signed-rank test with Benjamini-Hochberg correction (paired data) for non-normal distributions. For contingency tables, the pairwise Fisher’s exact test was used to compare each condition against a control, and *P*-values were adjusted for multiple comparisons using the Benjamini-Hochberg procedure.

### *In vivo* electroporation

*In vivo* electroporation of dCas9-KRAB mice was performed as described^11, 29^. The indicated plasmids were injected into the cerebellum of P6-8 mouse pups and then subjected to four 50 ms electrical pulses of 135 mV with 950 ms intervals. Electroporated pups were returned to moms and subjected to biochemical or immunohistochemical analyses at the indicated days after electroporation.

### Surgeries

Surgical procedures were performed as described with modifications^48^. Briefly, 8-18 weeks old mice were anesthetized with ketamine/xylazine (100/10 mg/kg intraperitoneal). Dexamethasone (2 mg/kg intramuscular), carprofen (5 mg/kg subcutaneous), and buprenorphine-SR (0.5-1.0 mg/kg subcutaneous) were administrated to minimize swelling of the brain and inflammation and to provide analgesia before the surgery. During surgery, lidocaine/epinephrine solution was applied to reduce bleeding and to provide local analgesia. After surgery, saline solution (37C, 0.3-0.5 ml subcutaneous) and carprofen (5 mg/kg subcutaneous) were given to replenish the animal’s fluids and to minimize inflammation, respectively. The head was shaved from the frontal bone to the occipital bone and eye ointment was applied. The surgical area was sterilized by wiping the skin with three alternating swipes of 70% ethanol and betadine.

For headplate and cannula implantation, animals were placed in a stereotaxic device (Stoelting) and the scalp overlaying the skull was removed. Under a surgical microscope, the fascia was removed, the skull surface was dried, and a custom head plate^10, 49^ was placed over the skull bregma and secured using Metabond cement (Parkell). A 0.8 mm hole in the skull over the anterior cerebellum was drilled and a mono fiber-optic cannula projecting 0.5 mm below the skull surface (0.22 NA, 200 µm core, Doric Lenses) was implanted. The cannula was secured to the skull using Metabond cement.

For viral delivery, a 0.8 mm hole in the skull was drilled over the anterior cerebellum. A fine glass pipette connected to a microinjector system (Nanoinject II, Drummond) and mounted in the stereotaxic frame (Stoelting) was used for injections. 0.5 µl of AAV (titer 1×10^13vg/ml) was injected at three different depths (0.2-0.4-0.7 mm) below the dura surface. The scalp was sutured and sealed with tissue glue (Vetbond). Experiments were conducted 3-5 weeks after viral infection.

### Open field

Mice at 11 weeks of age were placed in a rectangular cage (25 × 36 cm) and allowed to freely explore for 15 minutes, then transferred to a clean cage of the same dimensions for an additional 15 minutes. Following exploration, animals were returned to their home cages and sacrificed 2 hr later for immunohistochemistry.

### Inclined Treadmill

Mice at 11-17 weeks of age were habituated to a motorized treadmill (Maze Engineers, MZ-3.0-8-52) set at a 20° incline (3 minutes at 0 m/min, 10 minutes at 5 m/min) on the first day. On the following day, they underwent an exercise protocol consisting of 3 minutes at 0 m/min followed by six bouts of 5 minutes running, during which the speed was progressively increased from 0 to 18 m/min by 4 m/min each minute. Each bout was separated by 30 seconds at 0 m/min, for a total running duration of 30 minutes. Animals were returned to their home cages and sacrificed 10 min or 2 hr later for FANS-RNA-seq or protein immunohistochemical analyses, respectively.

### Delay Tactile Startle Conditioning

Delay Tactile Startle Conditioning (DTSC) was performed as described with modifications^10^ using littermate mice at 11-23 weeks of age. Following at least 5 days of recovery from surgery, head-fixed mice were habituated on a cylindrical treadmill for a 30 min session. For DTSC experiments in which trial initiation was coupled to locomotion, mice underwent an extended habituation consisting of 7-9 consecutive days of head fixation on the cylindrical treadmill. Habituation sessions lasted 15-40 min, with the duration gradually increased over successive days. Mice that failed to locomote reliably by the end of the habituation period were excluded from further experiments.

After habituation, mice underwent DTSC in a noise restricted room. Mice learned to make a rapid backward startle movement in response to an initially neutral conditioned stimulus (CS) consisting of a blue LED (Sparkfun) or optogenetic stimulation of granule neurons. The CS was paired with an unconditioned stimulus (US) consisting of a 30 ms aversive tactile stimulus delivered to the nose by an eagle plush toy (Amazon) secured to a motorized wheel (Sparkfun), with a CS-US inter-stimulus interval of 150 ms for the LED or 50 ms for the optogenetic stimulus. The shorter ISI for optogenetic stimulation elicits a conditioned response independently of visually induced conditioned responses, which require an ISI >100 ms^10, 49^. Mouse velocity and direction on the wheel were monitored using a rotary encoder, and trial onset was controlled by an Arduino Due board. Following a 15 s inter-trial interval, trials were initiated once the average running speed of the mouse over the preceding 3 s exceeded 5 cm/s, or after an additional 15 s if this criterion was not met.

Eighty DTSC trials, including CS-US paired trials and CS-only catch trials, were performed each day for up to ten days. Rotary encoder signals were recorded at 10 kHz using a Digidata 1440A Digitizer (Molecular Devices) synchronized to the visual or optogenetic CS and the tactile US.

For rest trials, trials with fewer than five rotary encoder transitions during the 100 ms before CS onset were included. Any forward or backward rotary encoder transition occurring 100-160 ms after LED CS onset or 0-60 ms after optogenetic CS onset was classified as a forward or backward conditioned response (CR), respectively. For locomotion trials, trials with a mean baseline velocity greater than 1 cm/s were included. An increase of more than 20% in maximum velocity or a decrease of more than 20% in minimum velocity between 20 and 80 ms after CS onset relative to the corresponding baseline value was classified as a forward or backward CR, respectively. The fraction or net fraction of backward CRs during each session was used as the index of sensorimotor learning.

### Optogenetic stimulation

A 450 nm diode blue laser (Opto Engine, 200 mW) coupled to a multimode fiber cable (Oz Optics) was used to deliver a 100 ms optogenetic stimulus composed of a 50 Hz train of 10 ms pulses and using 68 mW of laser power at the tip of the fiber to the ADCV of mice implanted with a fiber optic cannula.

For biochemical analyses, the ADCV of mice was subjected to 40 minutes of optostimulation (15 s inter-trial interval), after which the medial third of the ADCV was microdissected and processed for RNA-seq analyses.

### FANS

The dorsal cerebellum was collected from mice at 4, 8, 14, or 28 after electroporation. The anterior cerebellum was collected from control or transgenic mice at 3-9 weeks of age. The ADCV was collected from P16 mice or AAV infected mice at 12.5-14.5 weeks of age. Samples were fixed with homogenization in 1% PFA/PBS or 4% PFA/PBS solution, for RNA-seq or ChIP-seq and snRNA-seq, respectively, for 10 min, followed by quenching with glycine solution for 5 min. The cell pellet was washed twice with by BSA/PBS solution (0.3% BSA, 0.1% Triton-X in PBS) and stored at −80C until use. To isolate nuclei, the frozen pellet was thawed on ice, homogenized with lysis buffer (10 mM Tris-HCl pH 8, 10 mM NaCl, 0.2% Triton-X) and incubated on ice for 10 min. The lysate was pelleted by centrifugation at 700 *g* for 5 min at 4C and washed with lysis buffer once. The pellet was resuspended in BSA/PBS solution and filtered using a 50 µm filter (CellTrics). Filtered nuclei were incubated with the indicated primary antibodies, including mCherry (1:1000), GFP (1:5000), NeuN (1:500) or TUBB3 (1:1000), for 1 hr at 4C with rotation. Nuclei were washed with BSA/PBS solution twice and stained with secondary antibodies including anti-chicken Alexa 488 (1:250, Invitrogen, A11039), anti-mouse Alexa 647 (1:250, Thermo Fisher, A21235), anti-rabbit Alexa 488 (1:250, Thermo Fisher, A11034) or anti-rabbit Alexa 647 (1:250, Abcam, ab150075) for 30 min at 4C with rotation, followed by washing with BSA/PBS solution twice. Stained nuclei were resuspended in BSA/PBS solution and filtered using a 50 µm filter prior to FANS. Target nuclei were enriched with a SH800S cell sorter (Sony) using a 100 µm nozzle sorting chip and 488/638nm lasers. Nuclei were first sorted using the ultra-yield mode and then re-sorted using the normal mode (electroporated mice, and pSyt2-GFP and pNos1-GFP mice). Nuclei were sorted once using the ultra-purity mode (pNos(off)-mCherry mice). For NeuN sorting, target nuclei were enriched with BD FACSDiscover S8 Cell Sorter (BD Biosciences) using a 100 µm nozzle sorting chip and were sorted twice using the purity mode (P22 and P64 Cre-negative control mice). For snRNA-seq, NeuN^high^/TUBB3^low^ nuclei were sorted twice in the normal mode to increase purity. For RNA-seq, the lysis and incubation buffers contained 0.4 unit/μl of RNase inhibitor (Promega, N2515).

### RNA-seq

For optostimulation, the ADCV was homogenized in lysis buffer (50 mM Tris-HCl pH 8.0, 150 mM NaCl, 1% Triton-X, RNaseOUT (Thermo Fisher Scientific)) using a syringe and incubated on ice for 10 minutes. After spinning down, the supernatant was removed and the chromatin-associated RNA in the pellet was extracted with Trizol.

For FANS-RNA-seq, 2,000-4,500 sorted nuclei (Supplementary Table 5) from single or multiple animals were thawed on ice and reverse-crosslinked with 100 µL of RIP buffer (100 mM Tris-HCl pH 8.0, 10 mM EDTA, 1% SDS, 1 µl of RNAse inhibitor (Promega)) containing 1 µl of Proteinase K (New England Biolab) for 1 hr at 65C. Then, nuclear RNA was purified using a RNeasy micro kit (Qiagen) according to the manufacturer’s instructions.

5-7 ng of chromatin-associated RNA or half of the purified RNA was treated with a NEBNext rRNA Depletion Kit (New England Biolabs). RNA-seq was performed using libraries prepared with a NEBNext Ultra II Directional RNA Library Prep Kit for Illumina (New England Biolabs). All libraries were sequenced on the Illumina NextSeq 550 platform to obtain 37 bp paired-end reads. Two to eight biological RNA-seq replicates were performed.

### ChIP-seq

∼40,000-45,000 of sorted nuclei were subjected to sonication for DNA fragmentation in 0.5% SDS (Diagenode, Bioruptor pico) and the lysate were used for chromatin immunoprecipitation. Immunoprecipitation was performed in RIPA buffer (10 mM Tris-HCl pH 8.0, 140 mM NaCl, 0.1% SDS, 1% Triton-X, 0.1% DOC, 1 mM EDTA, 0.5 mM EGTA) using an antibody against H3K27ac with BSA-coated Dynabeads protein G (Thermo Fisher Scientific). After extensive washes of beads with RIPA buffer three times, high salt RIPA buffer three times, and TE three times, DNA fragments were eluted in elution buffer (10 mM Tris-HCl pH 8.0, 350 mM NaCl, 1% SDS, 0.1 mM EDTA at 65C for Dynabeads) for 30 min, treated with proteinase K for 1 hr at 37C, and de-crosslinked at 65C overnight. DNA fragments were purified with a PCR purification kit (Qiagen). Libraries were prepared using a NEBNext Ultra™ II DNA Library Prep Kit for Illumina (New England Biolabs) as per the manufacturer’s instructions. All libraries were sequenced on an Illumina NextSeq 550 platform to obtain 37 bp paired-end reads. Two to three biological ChIP-seq replicates were performed in all experiments.

### snRNA-seq

ADCV or the anterior lobe was microdissected from P16 C57BL/6 mice or from P49 or P52 B6D2F1/J mice, respectively, and fixed with homogenization in 4% PFA/PBS solution for 10min, followed by quenching with glycine solution for 5 min. Prior to library preparation, granule neuron nuclei were isolated by FANS using antibodies against NeuN and TUBB3, as described above. Sorted nuclei were fixed with crosslinking solution at 4C overnight and proceeded for library preparation according to the manufacturer’s protocol (Chromium Fixed RNA Profiling, 10x Genomics). Two biological replicates were performed.

### Immunohistochemistry

Cerebellar sections from adult mice 10-15 weeks of age, developmentally fate-traced mice at P24-P66, and electroporated mice at P14-P62 were prepared and labeled with the relevant antibodies as previously described and with Hoechst dye (Sigma B2261-25M)^26^. A modified heat-induced antigen retrieval protocol^50^ was used for ETV1 staining with sections were treated with a sodium citrate solution (10 mM sodium citrate, 0.05% Tween-20, pH 6.0) under high-pressure for 5 minutes using an Instant Pot® Duo Plus, followed by cooling to room temperature. Images were acquired at the Biological Imaging Facility (RRID:SCR_017767) on a confocal laser scanning microscope (Leica Microsystems, TCS SP8 confocal).

Granule neuron nuclei in the IGL of lobule IV/V were selected based on nuclear size and, when indicated, NeuN labeling. To calculate the IGL depth of granule neurons relative to the Purkinje cell layer (PCL) and white matter (WM), 10000 evenly spaced coordinates were interpolated along the PCL and WM boundaries and matched using mutual nearest neighbors to generate anchor points. Between each pair of anchors, additional point-to-point correspondences were interpolated to form a continuous series of smaller bounded regions that partitioned the IGL. Each cell centroid was assigned to its overlapping region, and its relative depth was calculated as the fractional distance from the PCL to the WM boundaries for that region.

For *in vivo* electroporation experiments, granule neurons in lobule IV/V and VIa were selected based on nuclear size and, when indicated, NeuN labeling. ETV1 and NOS1 immunofluorescence levels in mCherry-positive and NeuN-positive granule neurons subtracted by the median background intensity. ETV1 and NOS1 levels were then normalized to the background-subtracted median ETV1 intensity and 90th percentile NOS1 intensity, respectively, in mCherry-negative and NeuN-positive granule neurons.

For fate-tracing experiments, individual cells in the IGL were identified by segmenting LMNA-labeled nuclei using Cellpose-SAM (Cellpose 4)^51^. Only cells in the IGL were included, with granule neurons representing more than 95% of cells in this region^52, 53^. Segmented objects were included only if more than 30% of their pixels along the periphery exceeded the multi-Otsu threshold for LMNA fluorescence, their area was between 0.67 and 1.5 times the mean area of manually outlined nuclei, and their long-to-short axis ratio was no greater than 2.5. Mean YFP and NOS1 immunofluorescence intensities were measured within the segmented regions. For each image, intensities were background-subtracted using the mean intensity of cells below the 5th percentile and normalized to the background-subtracted mean intensity of cells above the 90th percentile. Cells were stratified into the YFP^high^, YFP^mid^, and YFP^low^ groups based on normalized intensity values of greater than 0.4, from 0.1 to 0.4, and below 0.1, respectively. Two to four mice were analyzed for all conditions.

### RNA-seq analyses

#### Alignment

Sequenced reads were aligned to the mm10 reference genome with HISAT2 using the public server at https://usegalaxy.org/ or https://usegalaxy.eu/ and normalized by library size. Gene annotations were derived using GENCODE version M11.

#### Differential gene expression and gene modules

Differential gene expression analyses were performed using likelihood-ratio tests for negative binomial generalized linear models with edgeR^54^ and the false discovery rate (FDR) was calculated for all genes.

For the *in vivo* electroporation time course, gene modules were generated from differentially expressed genes (FDR < 0.01) that were enriched at day 4 compared with day 8 (day 4/day 8 log_2_ fold change > 2), enriched at day 8 compared with both days 4 and 14 (day 8/day 4 and day 8/day 14 log_2_ fold changes >0.585, and cumulative log_2_ fold change >2), or enriched at day 14 compared with day 8 (day 14/day 8 log_2_ fold change >1). The resulting modules were visualized on the single-nucleus RNA-seq (snRNA-seq) UMAP plot.

Genetically-defined C1 (gC1) and C2 (gC2) gene modules were derived from a common set of differentially expressed transcripts (FDR < 0.01) identified by FANS-RNA-seq analyses of granule neuron subpopulations from pNos1-GFP, pSyt2-GFP, and pNos1(off)-mCherry mice. For comparisons of gene expression across late development, genes in these modules with |log_2_ fold change| > 1 between gC1 and gC2 were selected.

#### Mini-screen regression against the C1-C2 expression program

For each knockdown, linear regression was performed across the common set of genes used to define the gC1 and gC2 modules described above. For these genes, fold changes between control and knockdown conditions were regressed against the corresponding fold changes between GFP^high^ and GFP^low^ granule neurons in pSyt2-GFP mice, which represent the gC2 and gC1 subpopulations, respectively.

#### Gene Ontology

For gene ontology analyses, the common set of differentially expressed genes were retained if they showed the expected reciprocal expression pattern in the pNos1(off)-mCherry dataset relative to the pNos1-GFP and pSyt2-GFP datasets and had an absolute fold change > 1.5 between the gC1 and gC2 subpopulations in at least one dataset. Gene ontology analyses for genes enriched in gC1 or gC2 were performed against a background consisting of all expressed genes in the granule neuron FANS-RNA-seq datasets using the DAVID Bioinformatics Knowledgebase v2026_1^55^.

### snRNA-seq analyses

#### Alignment, dimensionality reduction, and clustering

snRNA-seq datasets were obtained from lobule IV/V granule neurons in P60 mice^12^ or generated from ADCV granule neurons in P16 mice or from anterior cerebellar granule neurons in P49 and P52 mice. For P16, P49, and P52 datasets, sequencing reads were demultiplexed and aligned to the mm10 transcriptome probe set using CellRanger 7.2, and gene expression matrices were generated. Using Seurat 4^56^, nuclei with more than 200 genes in the P60 datasets or more than 1000 genes in the P16, P49, or P52 datasets were retained for downstream analyses. This yielded a median of 1845-5522 genes per cell across datasets. Read counts were normalized, variance stabilized, and regressed for mitochondrial gene expression using sctransform, followed by dimensionality reduction with PCA preprocessing and UMAP embedding. Cell type annotation was performed iteratively through multiple rounds of dimensionality reduction and clustering^12^ to identify clusters enriched for granule neuron markers such as *Gabra6* or *Rbfox3* and remove cells enriched for cytoplasmic and mitochondrial genes.

For subsequent clustering of P60 granule neurons, Louvain clustering at a resolution of 0.1 identified the two major clusters, C1 and C2, and this partition resulted in the largest number of differentially expressed genes among the clustering resolutions tested. The same C1 and C2 clusters were obtained using Leiden clustering, along with a rare group of C3 cells marked by Chrm3. At higher resolutions, smaller clusters defined by more limited gene sets were detected. Granule neurons within the top quartile for expression of the top ten genes defining either the C1 or C2 cluster were selected to calculate average expression profiles for each subpopulation. Genes with log_2_ CPM >4 and |log_2_ fold change C2/C1| >0.585 were defined as C1 or C2 marker genes. Changes in these marker genes exhibited high reproducibility (correlation = 0.986) between two biological replicates.

For clustering of P16 granule neurons, Louvain or Leiden clustering at a resolution of 0.5 identified four major clusters. These clusters were defined by enriched marker genes, including developmentally regulated genes such as *Grin2b*, *Tmem91*, and *Snhg11*, which were previously characterized from bulk FANS-RNA-seq analyses^57^, and also independently defined using gene markers from *in vivo* electroporated granule neurons at day 4, day 8, and day 14.

#### Gene expression modules

Gene modules were derived from snRNA-seq clusters or from differential gene expression analyses of *in vivo* electroporated granule neurons. Module scores were calculated as the average expression of module genes in each nucleus, minus the average expression of control genes selected from expression bins matched to the population-averaged expression of module genes^56,58^.

For comparisons of C1 and C2 scores across datasets, scores were baseline-corrected by subtracting the bottom 1^st^ percentile of scores across cells within each dataset. The C1 and C2 identity of each granule neuron was determined by calculating the difference between the C1 and C2 scores, with neurons exhibiting higher C1 or C2 scores assigned to the corresponding identity. To assess the enrichment of *Etv1* mRNA among granule neurons subpopulations defined by the top, middle, or bottom third of C1-C2 scores, the fraction of granule neurons within each subpopulation whose *Etv1* expression ranked in the top 20% of all granule neurons was calculated.

### snATAC-seq analyses

#### Alignment, dimensionality reduction, and clustering

snATAC-seq datasets from granule neurons in P56-P63 mice including fragments and peaks were obtained^31, 32^. Using Seurat 4 with Signac^56, 59^, nuclei with more than 1000 fragments were retained for downstream analyses. Fragment counts were normalized using TF-IDF, and latent semantic indexing (LSI) was applied via SVD for dimensionality reduction prior to UMAP embedding. Gene activities were computed using fragments overlapping the gene body and the region 2 Kb upstream of the transcription start site (TSS). Clusters enriched for the granule neuron markers such as *Gabra6* or *Rbfox3* were retained for additional rounds of dimensionality reduction and clustering^56, 59^. Granule neuron snATAC-seq data was integrated with P60 granule neuron snRNA-seq data^12^ using canonical correlation analysis (CCA)^60^ to derive cell type labels for anterior or posterior granule neurons (projection score >0.5), which are enriched for *Rasgrf1*/*Ptprt* or *Gprin3*/*Galntl6*, respectively. Anterior granule neurons were then integrated with P60 lobule IV/V granule neuron snRNA-seq data^12^ using CCA to derive cell type labels for C1 or C2 granule neurons (projection score >0.5).

Differential snATAC-seq analyses for peaks overlapping with H3K27ac-marked enhancers were performed using likelihood-ratio tests based on negative binomial generalized linear models with EdgeR and calculating the FDR as described^10^. Transcription factor motif analyses were performed using Homer^61^ to identify known motifs within enhancer peaks that were enriched in C1 or C2 anterior granule neurons.

#### Chromatin accessibility modules

Chromatin accessibility modules were derived from differential enhancer analyses of granule neurons from pSyt2-GFP or dCas9-KRAB mice. Module scores were calculated as the average chromatin accessibility across enhancer peaks in each module per nucleus, minus the average accessibility of control peaks selected from bins matched to the population-averaged accessibility of module peaks^56, 58^. To quantify the transcriptional identity of granule neurons, the GFP^low^ score minus GFP^high^ score was calculated for each nucleus.

### ChIP-seq analyses

ChIP-seq reads were aligned to the mm10 reference genome with Bowtie2 using the public server at https://usegalaxy.org/ or https://usegalaxy.eu/ and normalized by library size. Peak calling was performed using MACS2^62^ and those overlapping the ENCODE blacklist were removed^63^. H3K27ac peaks >2 kb distal to H3K4me3-marked TSSs were considered as enhancers.

Differential H3K27ac ChIP-seq peak analyses and transcription factor motif analyses were performed as described above for snATAC-seq data to identify known motifs located within enhancers enriched in GFP^low^ and GFP^high^ granule neurons from pSyt2-GFP mice, or within enhancers showing differential activity in control and knockdown granule neurons from dCas9-KRAB mice.

For regression analyses, a common set of differentially active enhancers from FANS-H3K27ac ChIP-seq analyses of granule neuron subpopulations in pNos1-GFP (FDR < 0.05) and pSyt2-GFP (FDR < 0.1) was curated. For these enhancers, pairwise linear regression analyses were performed to compare fold changes in H3K27ac levels between control and knockdown conditions with fold changes between the GFP^high^ and GFP^low^ granule neurons in pSyt2-GFP mice, representing the genetically-defined C2 and C1 subpopulations, respectively.

## Notes

### Competing Interest Statement

The authors have declared no competing interest.

