## Supplementary information for "Postmitotic Regulation of Neuronal Subpopulations in the Cerebellum"

### Extended Data Figure Legends

**Extended Data Fig. 1. Differential gene expression in C1 and C2 granule neurons.**

**a**, **b**, Left: expression of *Syt1* and *Cadm1* (**a**) or *Cntnap4* (**b**) among granule neurons in the snRNA-seq UMAP from Fig. 1b. The $\log_2$ fold change in expression between C1 versus C2 granule neuron subpopulations is indicated (C1-C2). Right: expression of these C1 (**a**) or C2 (**b**) enriched genes in Allen Brain Atlas *in situ* hybridization images of the internal granule layer of cerebellar lobule IV/V. ML: molecular layer, PCL: Purkinje cell layer, IGL: internal granule layer, WM: white matter. **c**, Fluorescence-activated nuclei sorting (FANS) procedure for isolating granule neurons using antibodies targeting mCherry and NeuN. The fluorescence intensities of labeled cells and the sorting gates are shown for mCherry$^{high}$/NeuN$^{high}$ (red rectangle) and mCherry$^{low}$/NeuN$^{high}$ (red circle) populations from pNos1(off)-mCherry mice. **d**, Comparison of $\log_2$ fold changes in gene expression between GFP$^{high}$ and GFP$^{low}$ populations in pNos1-GFP mice (left) or between mCherry$^{high}$ and mCherry$^{low}$ populations in pNos1(off)-mCherry mice (right) with $\log_2$ fold changes in gene expression between C1 and C2 clusters from snRNA-seq analyses. The correlation coefficients (correl) are indicated. **e**, **f**, Expression of the top 100 C1 or C2 marker genes (**e**, two-sided Wilcoxon signed-rank test) or $\log_2$ fold change (FC) in immediate-early gene expression (IEG, **f**; two-sided one-sample *t*-test against 0, $n$ = 4 biological replicates) in FANS-isolated granule neurons from the mouse lines shown in Fig. 1e, g, h. FP: fluorescent protein. **g**, Expression of *Fos* (left) and *Npas4* (right) among granule neurons in the snRNA-seq UMAP from Fig. 1b.

Box plots in **e** show median, quartiles (box), and range (whiskers). Data in **f** show mean and error bars denote s.e.m. *$P$ < 0.05, ** $P$ < 0.01, ***$P$ < 0.001.

**Extended Data Fig. 2. Gene expression in developing granule neurons.**

**a**, FANS procedure for isolating granule neuron nuclei using antibodies against the granule neuron marker NeuN and the cytoplasmic protein TUBB3. Fluorescence intensities of labeled cells and sorting gates are shown (box). **b**, Expression of the *Grin2b*, *Tmem91*, and *Snhg11* marker genes corresponding to different stages of granule neuron development, visualized on the snRNA-seq UMAP from P16 cerebellum in Fig. 2b. **c**, UCSC genome browser tracks showing *Grin2b* and *Snhg11* expression in developing granule neurons isolated by FANS after *in vivo* electroporation. **d**, Distribution of C1 and C2 gene scores among anterior cerebellar granule neurons from snRNA-seq analyses of P16 and P60 mice.

**Extended Data Fig. 3. Late developmental changes in granule neuron gene expression.**

**a**, Distribution of C1 and C2 gene scores in granule neurons as in Extended Data Fig. 2d stratified into the bottom, middle, and top thirds by *Rbfox3* levels (two-sided Wilcoxon rank-sum tests, $n$ = 6027, 6026, 6026 cells for bot, mid, top at P16 and 57758 cells at P60). **b**, Labeled regions in lobule IV/V after electroporation at P14, P20, P34, and P62, with dashed lines indicating the IGL boundary as in Fig. 3f. **c**, Tamoxifen injection strategy in juvenile P17 or adult P56 transgenic mice to fate-label granule neurons with ChR2-YFP in a *Nos1*-dependent manner, as in Fig. 3h. Seven weeks after injections, mice were subjected to immunohistochemistry with antibodies against YFP and LMNA, the latter labeling nuclei. **d**, Distribution of YFP levels in IGL cells (two-sided Wilcoxon rank-sum test, $n$ = 19550, 7562 cells from 3 juveniles, 2 adults).

Box plots in **a**, **d** show median, quartiles (box), and range (whiskers). ***$P < 0.001$.

**Extended Data Fig. 4. Genetic mini-screen for regulators of C1 and C2 granule neurons**

**a**, Fold change in mRNA levels upon knockdown (KD) of mini-screen targets relative to their respective control conditions (two-sided one-sample *t*-test against 1, from left to right, $n$ = 6, 6, 7, 2, 6, 3, 6, 8, 2, 6, 6, 6, 6, 6 biological replicates). **b**, Overlap between genes downregulated upon ETV1 knockdown (FDR < 0.01, two-sided $P$ value from a likelihood-ratio test using a negative binomial generalized linear model with Benjamini-Hochberg correction, $n$ = 6 biological replicates) and C1 or C2 marker genes, with the Jaccard index indicating the extent of overlap. **c**, **d**, Jaccard index (**c**) and the overlap count (**d**) for genes downregulated upon knockdown or knockout of mini-screen targets (FDR < 0.01, two-sided $P$ value from a likelihood-ratio test using a negative binomial generalized linear model with Benjamini-Hochberg correction, $n$ as in **a**) and C1 or C2 marker genes. **e**, Expression scores of genes downregulated by ETV1 knockdown in $C1^{high}$ and $C2^{high}$ granule neurons at P16, P49, P52, and P60, as in Fig. 2e (two-sided *t*-test, from left to right, $n$ = 5453, 12625, 5100, 6437, 84420, 88827 cells). **f**, **g**, Expression scores of genes downregulated by GRIN1 knockdown (**f**) or CRHR1 knockdown (**g**) compared to control neurons, overlaid onto the snRNA-seq UMAP of P16 granule neurons as in Fig. 2f. **h**, UCSC genome browser tracks showing expression of the C2 marker gene *Syt2* in control granule neurons, or following ETV1 knockdown or combined GRIN1 and CACNA1C knockdown. **i**, **j**, $Log_2$ fold change in *Syt2* (**i**), *Kcnip4* (**j**, left) or *Plekhg3* (**j**, right) in ETV1 knockdown, GRIN1 knockdown, CACNA1C knockdown, or the combined GRIN1 and CACNA1C knockdown condition, relative to the control condition (from left to right, $n$ = 6, 6, 6, 4 biological replicates). **k**, Regression coefficients showing the effects of GRIN1 knockdown, CACNA1C knockdown, or combined GRIN1 and CACNA1C knockdown on the expression of genes downregulated by ETV1 knockdown.

Box plots in **e** show median, quartiles (box), and range (whiskers). Data in **a**, **i**, and **j** show mean and error bars denote s.e.m. $*P < 0.05$, $** P < 0.01$, $***P < 0.001$.

**Extended Data Fig. 5. Roles of ETV1 in chromatin accessibility and its regulation by calcium signaling**

**a**, UCSC genome browser tracks showing gene expression and H3K27ac levels in genetically-defined $GFP^{low}$ (gC1) and $GFP^{high}$ (gC2) granule neurons from pSyt2-GFP mice at the *Clmp*, *Map4k4*, and *Tmem132c* loci, which represent a C1 marker gene, a gene shared by C1 and C2, and a C2 marker gene, respectively. Orange and cyan arrows denote enhancers. **b**, Relative H3K27ac enrichment in gC1 and gC2 neurons at enhancers, as in **a**, that overlap genes with enriched expression in $GFP^{low}$ (gC1) or $GFP^{high}$ (gC2) granule neurons from pSyt2-GFP mice, as in Fig. 1k, or genes with expression shared between subpopulations (two-sided Wilcoxon rank-sum tests against the shared genes condition with Benjamini-Hochberg correction, $n$ = 88, 710, 56 loci for gC1, shared, gC2). **c**, Transcription factor binding motifs enriched in accessible enhancer regions that are preferentially active in C1 versus C2 granule neurons, as identified from integrated snATAC-seq and snRNA-seq analyses. **d**, Chromatin accessibility scores at enhancers downregulated by ETV1 knockdown compared to control neurons, overlaid onto the snATAC-seq UMAP in Fig. 5a. **e**, Images of control (ctrl) or combined GRIN1 and CACNA1C knockdown granule neurons labeled for ETV1 (gray) and mCherry (red), together with the Hoechst DNA dye (blue). Dashed lines indicate nuclear boundaries of mCherry-labeled granule neurons. **f**, Quantification of ETV1 immunofluorescence levels in granule neurons as shown in **e** (two-sided one-sample *t*-test against 1, from right to left, $n$ = 2, 3, 3, 3 biological replicates). **g**, Fold change in *Etv1* mRNA levels in GRIN1 knockdown, CACNA1C knockdown, or combined GRIN1 and

CACNA1C knockdown granule neurons compared to control neurons (from right to left: $P = 1.4 \times 10^{-2}$, $7.9 \times 10^{-3}$, 0.10; two-sided one-sample *t*-test against 1, $n$ = 6, 6, 4 biological replicates). Box plots in **a** show median, quartiles (box), and range (whiskers). Data in **f** and **g** show mean and error bars denote s.e.m. *$P < 0.05$, ***$P < 0.001$.

**Extended Data Fig. 6. Differential activity of granule neurons with sensorimotor experience**

**a**, Schematics of the home cage condition and inclined treadmill paradigm. **b**, Images of ADCV sections infected with AAV9-pNpas4-shGFP and labeled for GFP (green) and Hoechst (blue) from mice subjected to the behavioral tasks shown in **a**. Arrows indicate GFP-positive active cells in the IGL. **c**, **d**, Quantification of the number of GFP-positive cells within the field of view (**c**, one-way ANOVA with Dunnett's post hoc test against the home cage condition, $n$ = 6, 10, 6 regions in 3, 3, 4 mice for homecage, open field, inclined treadmill) or their location relative to the PCL and WM (**d**, analyzed as in Fig. 1i; two-sided Wilcoxon rank-sum tests against the home cage condition with Benjamini-Hochberg correction, $n$ = 54, 214, 356 cells for homecage, open field, inclined treadmill) in mice subjected to the home cage condition, open field paradigm, or the inclined treadmill paradigm. **e**, mRNA expression of *Fos* and *Npas4* in $\text{mCherry}^{\text{high}}$ and $\text{mCherry}^{\text{low}}$ granule neurons populations, corresponding to C1 and C2 granule neurons, respectively, in pNos1(off)-mCherry mice subjected to the home cage condition ($\log_2$FC shown in Extended Data Fig. 1f) or the inclined treadmill paradigm (two-sided *t*-test, $n$ = 4, 3 mice for homecage, inclined treadmill).

Box plots in **d** show median, quartiles (box), and range (whiskers). Data in **c**, and **e** show mean and error bars denote s.e.m. *$P < 0.05$, **$P < 0.01$, ***$P < 0.001$.

**Extended Data Fig. 7. Differential activity of granule neurons with sensorimotor experience**

**a**, Images of ADCV granule neurons in the IGL from GN-ChR2-YFP or $GN^{C2}$-ChR2-YFP mice labeled for GFP/YFP (green) and Hoechst (gray) as in Fig. 6b. **b**, The fraction of YFP-positive granule neurons among all cells in the IGL for the conditions shown in **a** (*n* = 2, 4 biological replicates for GN-ChR2-YFP, $GN^{C2}$-ChR2-YFP). **c**, **d**, mRNA levels of *Fos* (**c**) or *Npas4* (**d**) induced by optostimulation in the ADCV of GN-ChR2-YFP or $GN^{C2}$-ChR2-YFP mice ($log_2FC$ shown in Fig. 6c; two-sided *t*-test, from left to right, *n* = 6, 7, 5, 4 biological replicates). **e**, Schematic of the delay tactile startle conditioning paradigm using a blue LED as the CS paired with a tactile stimulus to the nose as the US. **f**, **g**, Mice subjected to multiple days of conditioning using the protocol in **e**. CR percentages (**f**, *n* = 9 mice) and traces during catch trials (**g**, *n* = 6 mice) are shown. A backward CR to the LED CS was acquired over days, with a 100 ms delay after CS onset, consistent with previous observations using visual stimuli in cerebellar conditioning[10, 38]. Data in **c**, **d**, **f**, and **g** show mean and error bars denote s.e.m. $^{**}P < 0.01$, $^{***}P < 0.001$.

**Supplementary Tables**

Supplementary Table 1. C1 and C2 marker genes from snRNA-seq and FANS-RNA-seq analyses.

Supplementary Table 2. Differentially expressed genes (FDR < 0.01) in granule neurons across days 4, 8, and 14 after electroporation.

Supplementary Table 3. Mouse lines and samples used in this study.

Supplementary Table 4. Sequences of sgRNAs for the *in vivo* genetic mini-screen.

Supplementary Table 5. Sample information for RNA-seq analyses.

**Extended Data Fig. 1**

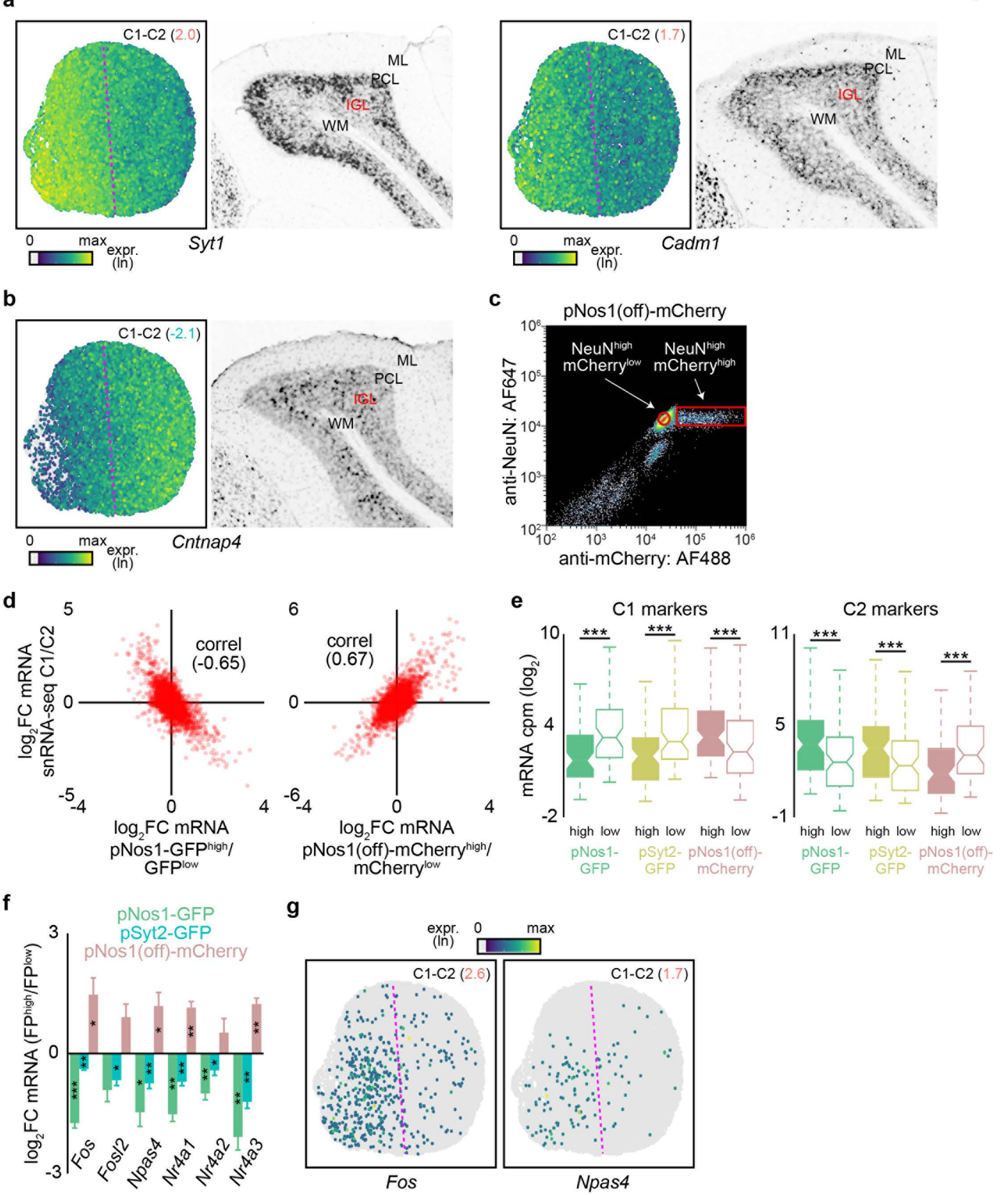

### Extended Data Fig. 2

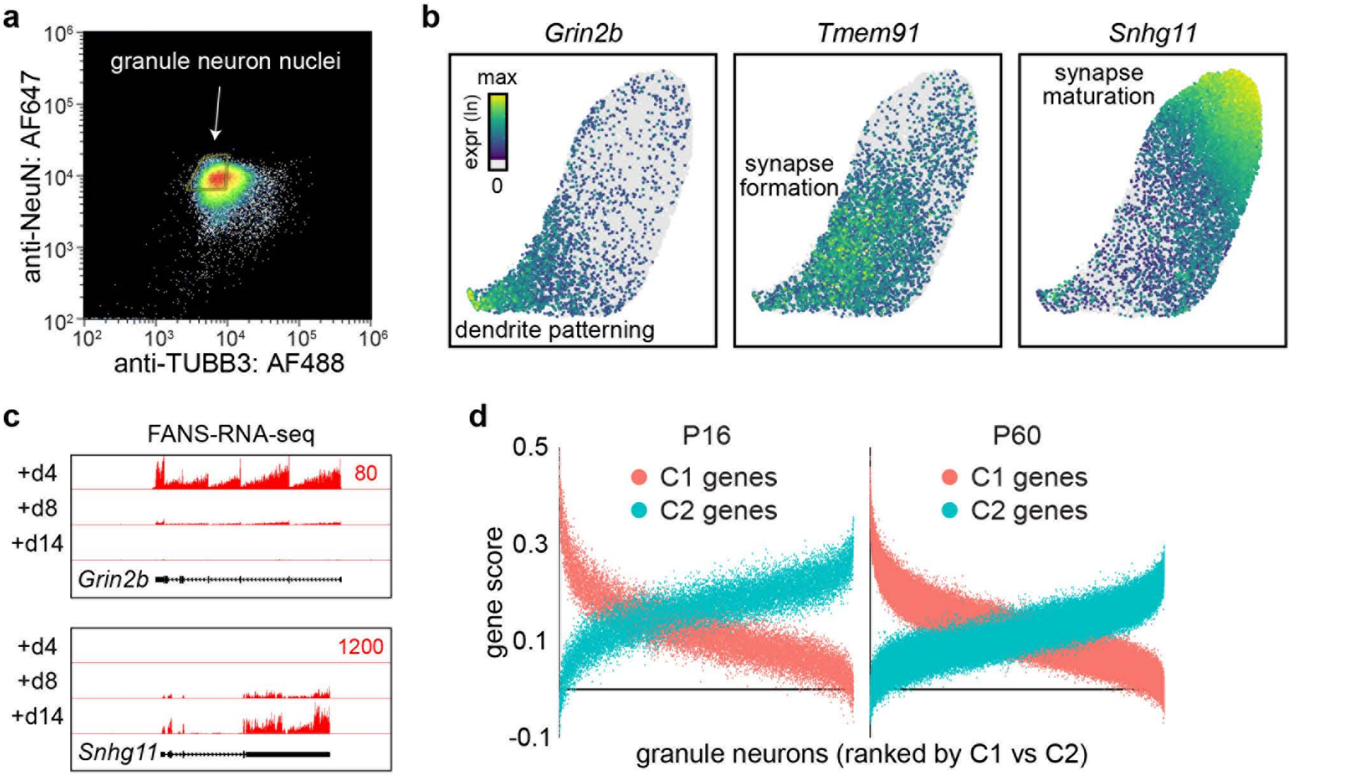

**Extended Data Fig. 3**

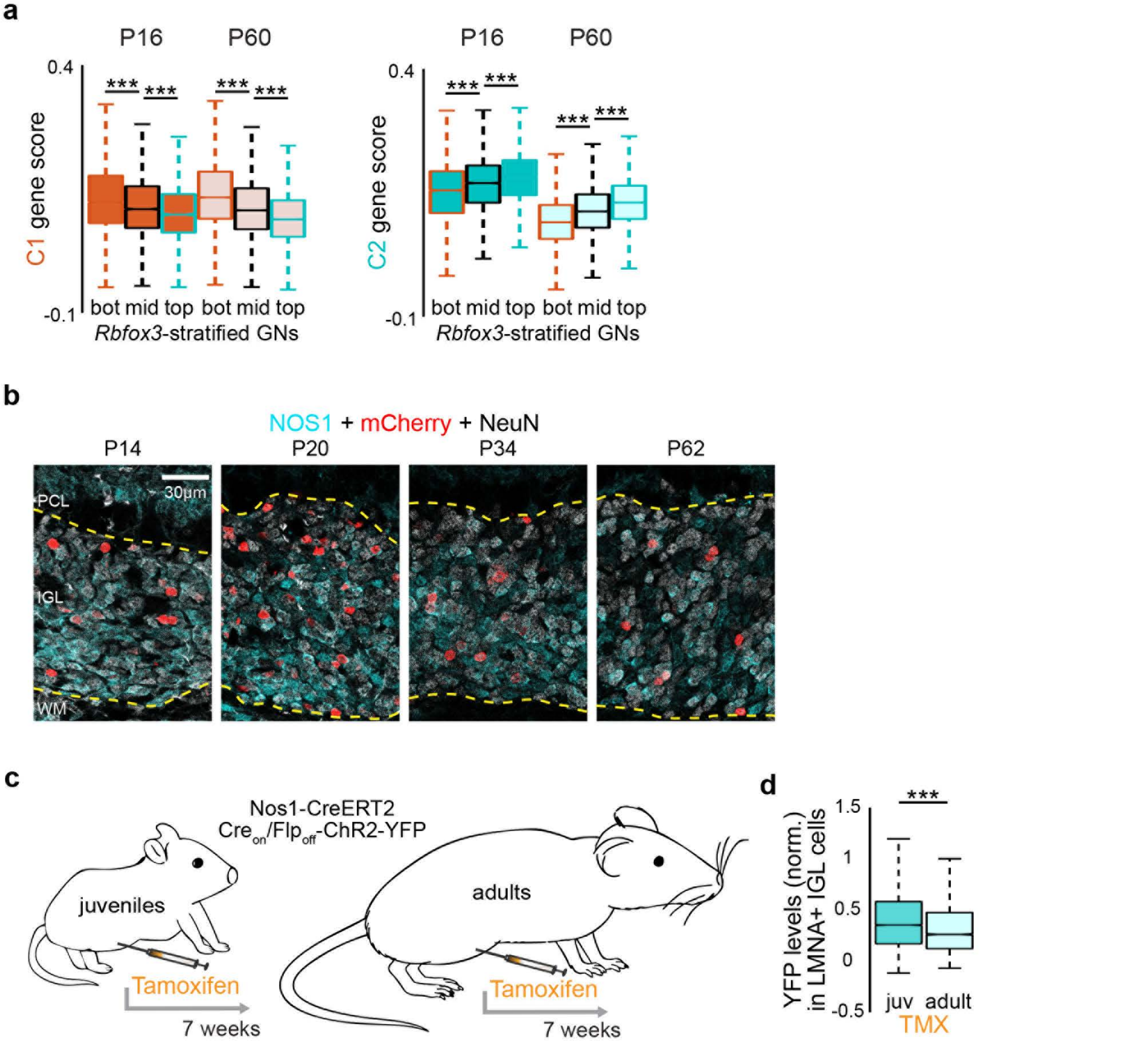

Extended Data Fig. 4

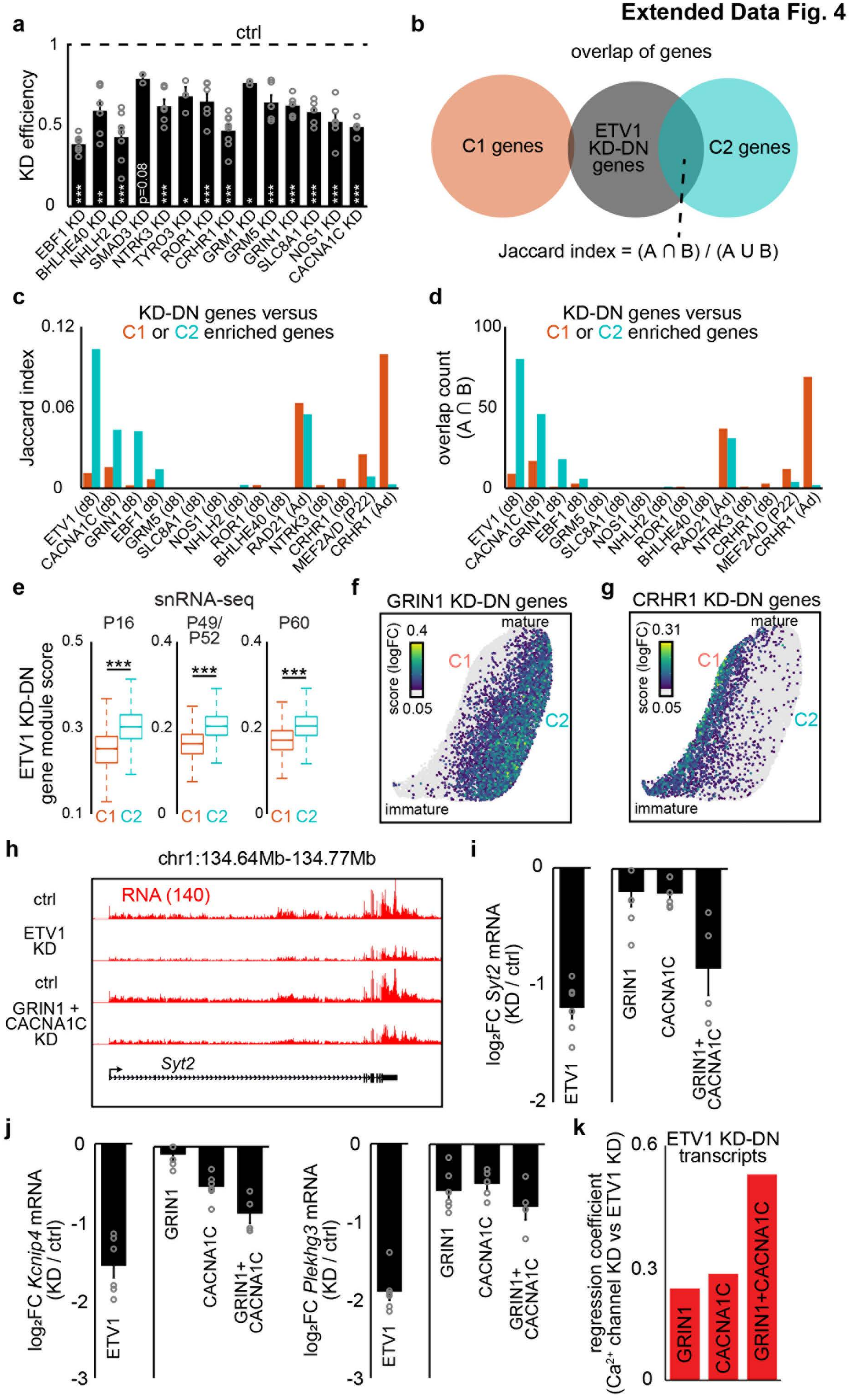

### Extended Data Fig. 5

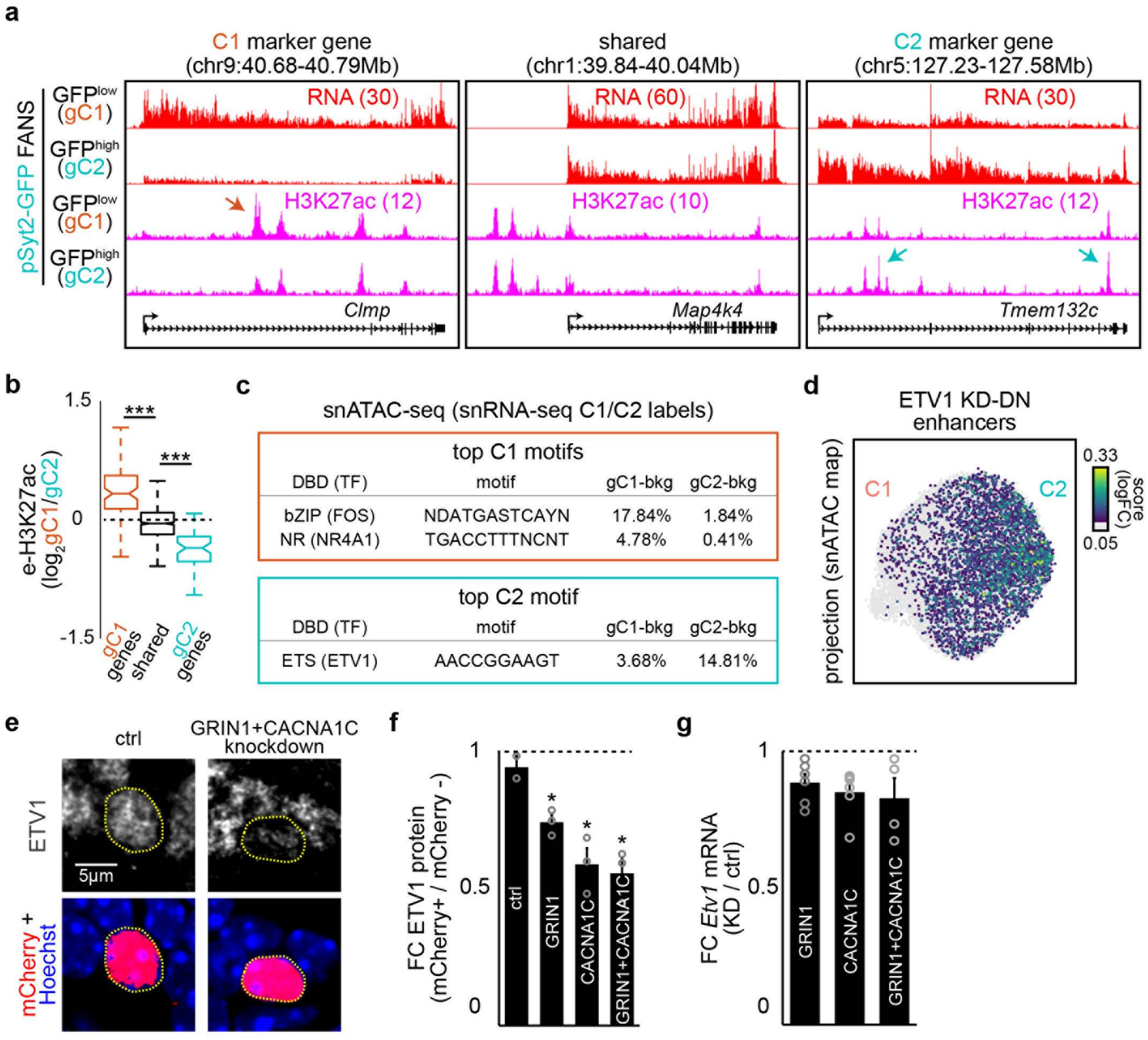

### Extended Data Fig. 6

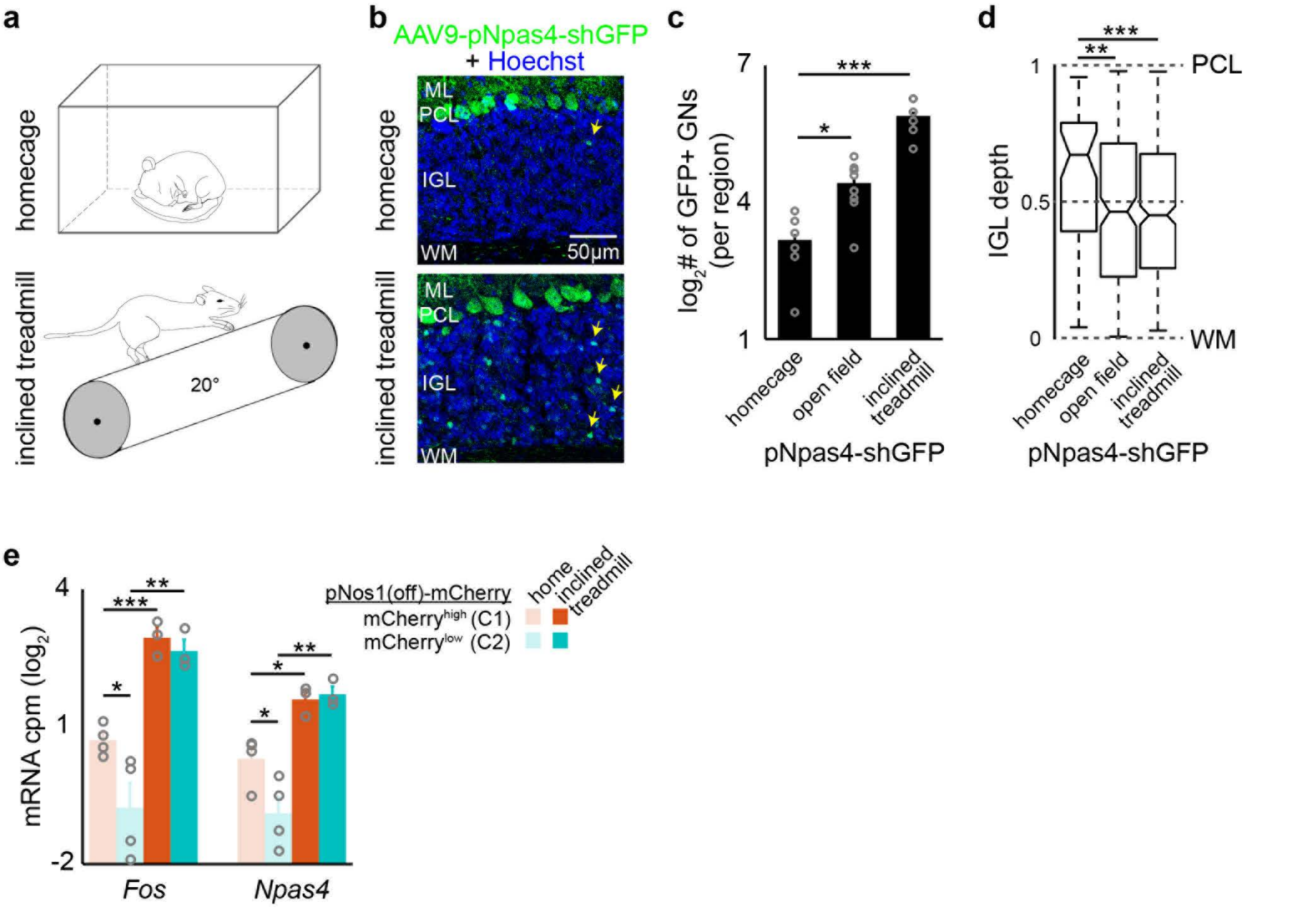

**Extended Data Fig. 7**

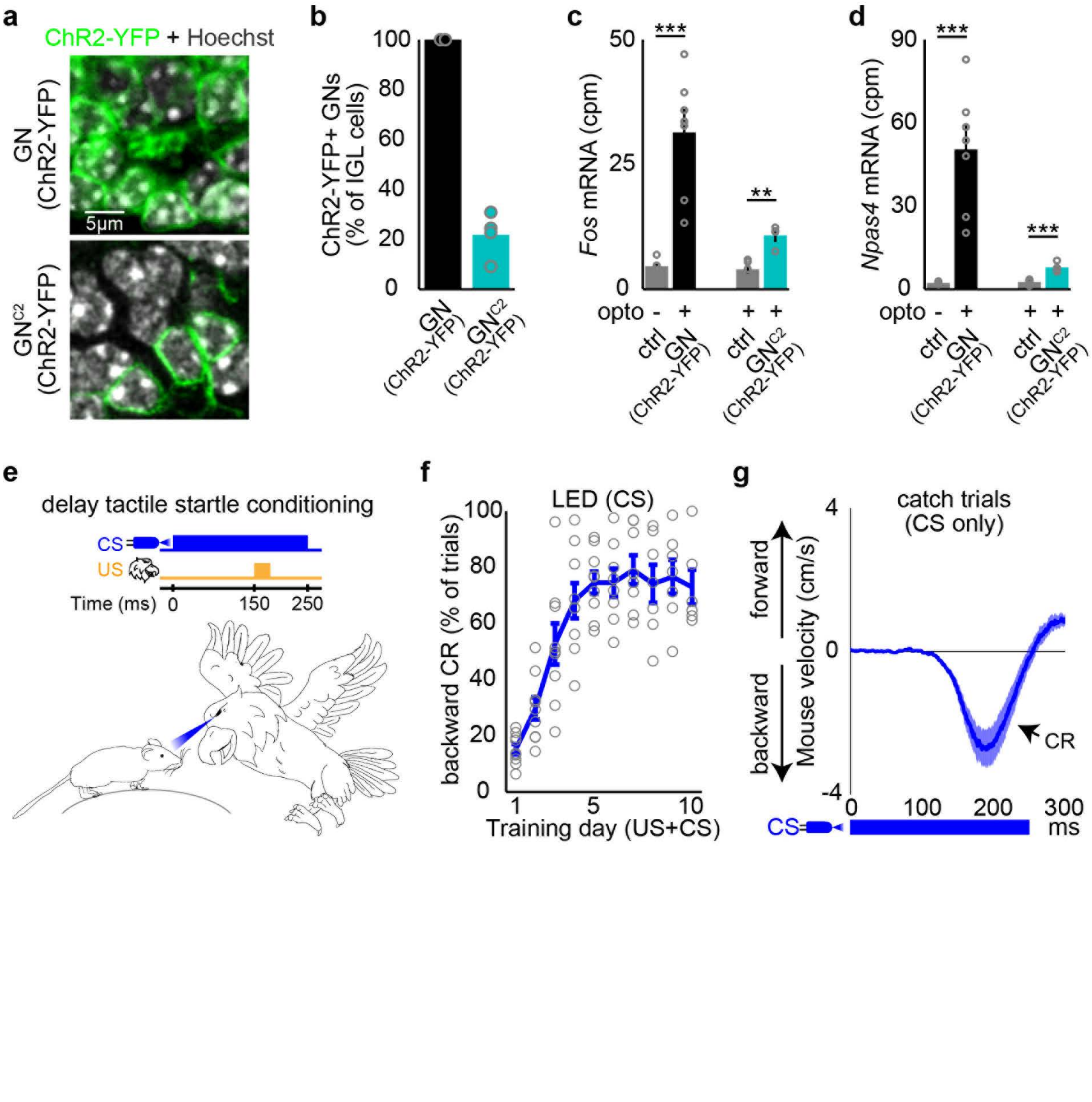
